# Modulation of the agonist and antagonist activity of peptidic FPR1 ligands through N-terminal modifications: A structural and functional analysis

**DOI:** 10.64898/2026.07.10.737841

**Authors:** Sarah Maskri, Denise Pajonczyk, Joana Massa, Carsten Raabe, Thomas Bödeker, Bernhard Wünsch, Ursula Rescher, Oliver Koch

## Abstract

Formyl peptide receptor 1 (FPR1) is a promising therapeutic target for the treatment of inflammatory and infectious diseases. Although multiple classes of peptides are known to modulate FPR1 activity, comprehensive studies systematically linking N-terminal modifications to binding, mechanism of action and functional outcomes remain limited. In this study, we aimed to rationalise the binding and activity of three peptide series (MLF, FLFLF, and MLFYLA) featuring diverse N-terminal modifications from a structural point of view. A combined *in silico* and *in vitro* approach was employed to evaluate the activity of newly designed peptide agonists and to generate mechanistic binding hypotheses. Our findings led to the identification of a transmembrane binding pocket in FPR1, which provides a structural basis for the observed antagonist and partial agonist behaviours and leads to a generalisable strategy for tuning the functional outcome of peptidic ligands.

## Introduction

G protein-coupled receptors (GPCRs) constitute the most important target class in modern drug discovery.^[1,2]^ These membrane proteins share a conserved seven-transmembrane-helical architecture and regulate numerous physiological processes by transducing signals through heterotrimeric G-proteins and other intracellular effectors. Dysregulation of GPCR signalling is implicated in a wide range of diseases, highlighting their considerable therapeutic relevance.^[3–5]^

The human formyl peptide receptor (FPR) belongs to class A GPCR that sense pathogen- and danger-associated molecular patterns (PAMPs and DAMPs) and comprises three members (FPR1, FPR2, and FPR3).^[6–8]^ More specific, FPRs recognize N-formylated peptides derived from bacteria or mitochondria. In bacteria, protein synthesis is initiated with N-formyl methionine, a modification absent from eukaryotic proteins, making N-formylated peptides reliable markers of bacterial presence. Because mitochondria share a bacterial evolutionary origin, mitochondrially encoded proteins are similarly N-formylated, enabling damaged cells to release these peptides as DAMPs.^[9–12]^

FPR ligands are structurally diverse, ranging from short N-formylated peptides such as the prototypical agonist fMLF to longer peptides and synthetic analogues like the hexapeptide agonist WKYMVm (W-peptide), identified through combinatorial peptide library screening.^[9,13]^ Structural studies show that FPR1 and FPR2 share ∼69% amino acid sequence identity overall, with particularly strong conservation in the intracellular region (>90%). Especially many ligand-binding residues are conserved, suggesting related recognition mechanisms while allowing selectivity.^[14–16]^

At the molecular level, recognition of formylated peptides is mediated by a conserved polar triad consisting of residues D106^3.33^, R201^5.38^, and R205^5.42^ in FPR1. Mutagenesis studies have demonstrated that substitution of any of these residues strongly reduces agonist activity, highlighting their central role in ligand binding and receptor activation. The polar triad is thought to stabilise the insertion of hydrophobic side chains such as methionine into the “activation chamber” formed by L109^3.36^, F110^3.37^, V113^3.40^, and W254^6.48,^ a hydrophobic pocket at the base of the orthosteric site.^[15,17]^

Within this family, FPR3 is largely insensitive to N-formylated peptides due to differences in key binding-site residues.^[14]^ FPR2 can recognize formylated peptides but is less excitable, whereas FPR1 exhibits high sensitivity, with N-formylated peptides often acting as superagonists.^[8,10,18,19]^ These properties establish FPR1 as the primary sensor of bacterial and mitochondrial danger signals and rendering FPR1 antagonism an attractive strategy in infection and inflammation control.^[20,21]^

Interestingly, small N-terminal modifications of peptide ligands can convert FPR1 agonists into antagonists. Replacement of the N-terminal formyl group in fMLF with a tert-butyloxycarbonyl (Boc) protecting group produces the antagonist Boc-MLF. Similarly, Boc-FLFLF represents another well-characterised FPR1 antagonist.^[22,23]^ The molecular basis of this agonist-to-antagonist switch was not fully understood, particularly regarding how subtle N-terminal modifications alter receptor activation while maintaining the same peptide sequence. We therefore carried out a comprehensive computational analysis in combination with pharmacological characterisation in order to gain a better understanding of the effects of these modifications.

**Figure 1.**
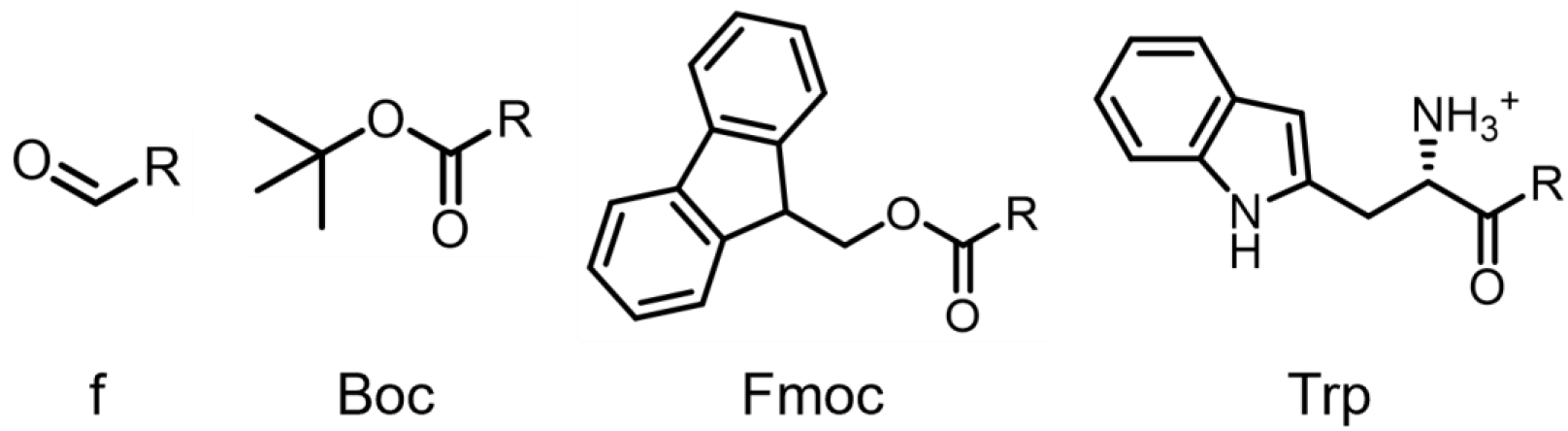
Structures of the four N-terminal modifications applied to three peptide series in this work.

We used homology modelling, molecular dynamics simulations, and *in vitro* pharmacological assays to investigate the structural determinants that govern agonism and antagonism at FPR1. Using the known agonist–antagonist peptide pair fMLF/Boc-MLF, we explored how N-terminal hydrophobic modifications can tune agonistic and antagonistic properties. This analysis was further extended to novel N-terminal modifications such as fluorenylmethoxycarbonyl (Fmoc) and the introduction of the additional amino acid tryptophan (Trp). A third peptide series derived from SP8 (fMLFYLA) was included to assess the generality of these structural determinants.

Collectively, this study provides mechanistic insights into ligand binding at FPR1 and identifies molecular determinants governing the transition from agonism to antagonism. These findings enhance understanding of FPR1 ligand recognition and provide a framework for the structure-based design of modulators targeting inflammatory and immune-related diseases.

## Results

At the beginning of this project, a FPR1 protein structure was not available. Therefore, an FPR1 homology model was generated based on the active-state FPR2 crystal structure in complex with WKYMVm (PDB ID: 6LW5^[16]^). The model was further refined by three independent 1 μs molecular dynamics (MD) simulations to enable structural relaxation and improve model quality. The resulting structure of these simulations (Figure 2A) shows good agreement with the later reported cryo-EM structures of FPR1 bound to formylated peptides (PDB IDs 7EUO, 7T6T, 7VFX and 7WVU; SI Table S1), supporting its reliability for pose generation.^[15,17,24]^

**Figure 2.**
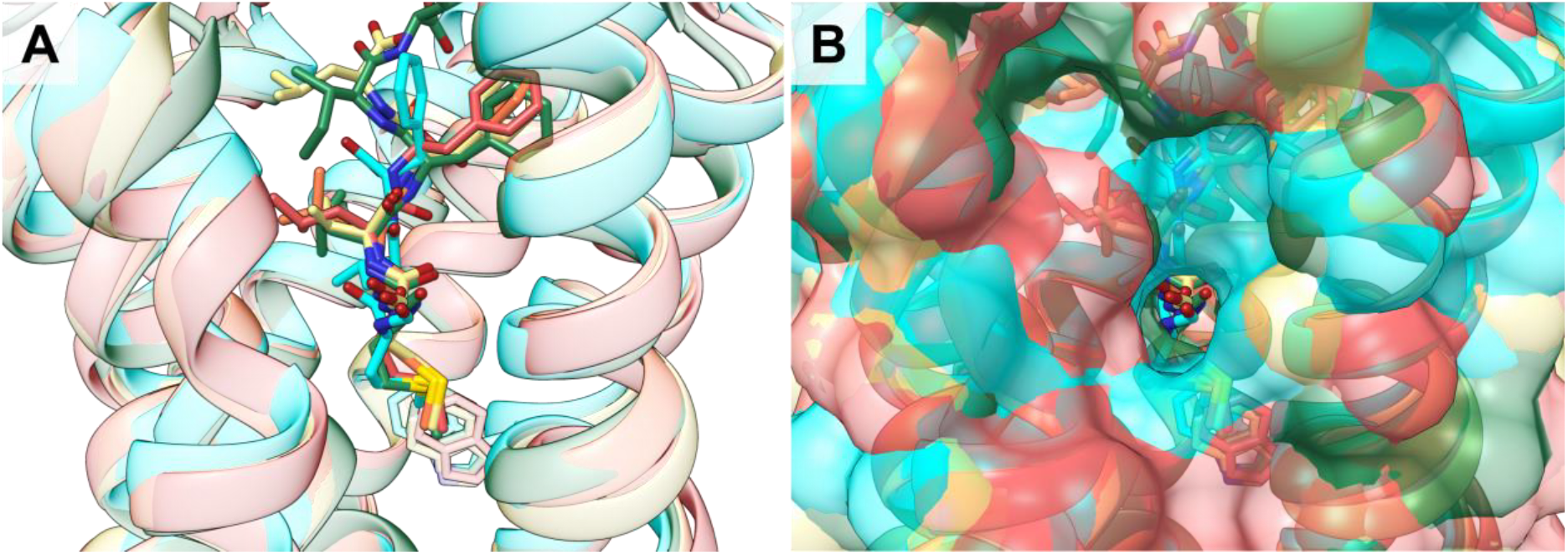
A) The validated homology model of FPR1 and the binding pose of fMLF. Four experimental structures of FPR1 are available which are all bound to formylated peptides with the PDB IDs 7EUO^[17]^, 7T6T^[15]^, 7VFX^[17]^ and 7WVU^[24]^ (coloured red, orange, yellow and green, respectively). These structures are aligned with the modelled pose of fMLF (cyan). B) Cavity visible between TM3, TM4 and TM5 in the homology model and experimental structures of FPR1. It is lined by the residues F110^3.37^, V113^3.40^ and I204^5.41^ and R205^5.42^.

### Binding Modes of fMLF and Boc-MLF reveal determinants of FPR1 activation and antagonism

FPRs can bind various peptides that have been isolated from bacteria. Commonly, these peptides feature a formylated N-terminal methionine, suggesting that this moiety is essential for binding to FPR1 and FPR2.^[25]^ The chemoattractant fMLF found in E. coli is a potent and quite selective FPR1 agonist that binds with its formylated N-terminus inserted deep into the orthosteric site and its C-terminus pointing outward.^[15,17,24]^ To investigate the molecular determinants of FPR1 activation, we first analysed the binding mode of the canonical orthosteric agonist fMLF. Docking studies and subsequent MD simulations revealed a hydrogen (H-) bond network involving the polar triad D106^3.33^, R201^5.38^, and R205^5.42^ extended by interactions with Y257^6.51^. The tight interactions by the formyl further enables the methionine of the peptide to be inserted into a deep hydrophobic cavity (“activation chamber”), formed by L109^3.36^, F110^3.37^, V113^3.40^, and W254^6.48^. The negatively charged C-terminus of fMLF is stabilised by electrostatic interactions with either R84^2.63^ or K85^2.64^ (Figure 3A). The lack of this C-terminal stabilising interactions might explain fMLF’s preference for FPR1 over FPR2, which is consistent with the reduced activity observed for the R84^2.63^D mutant.^[24]^

**Figure 3.**
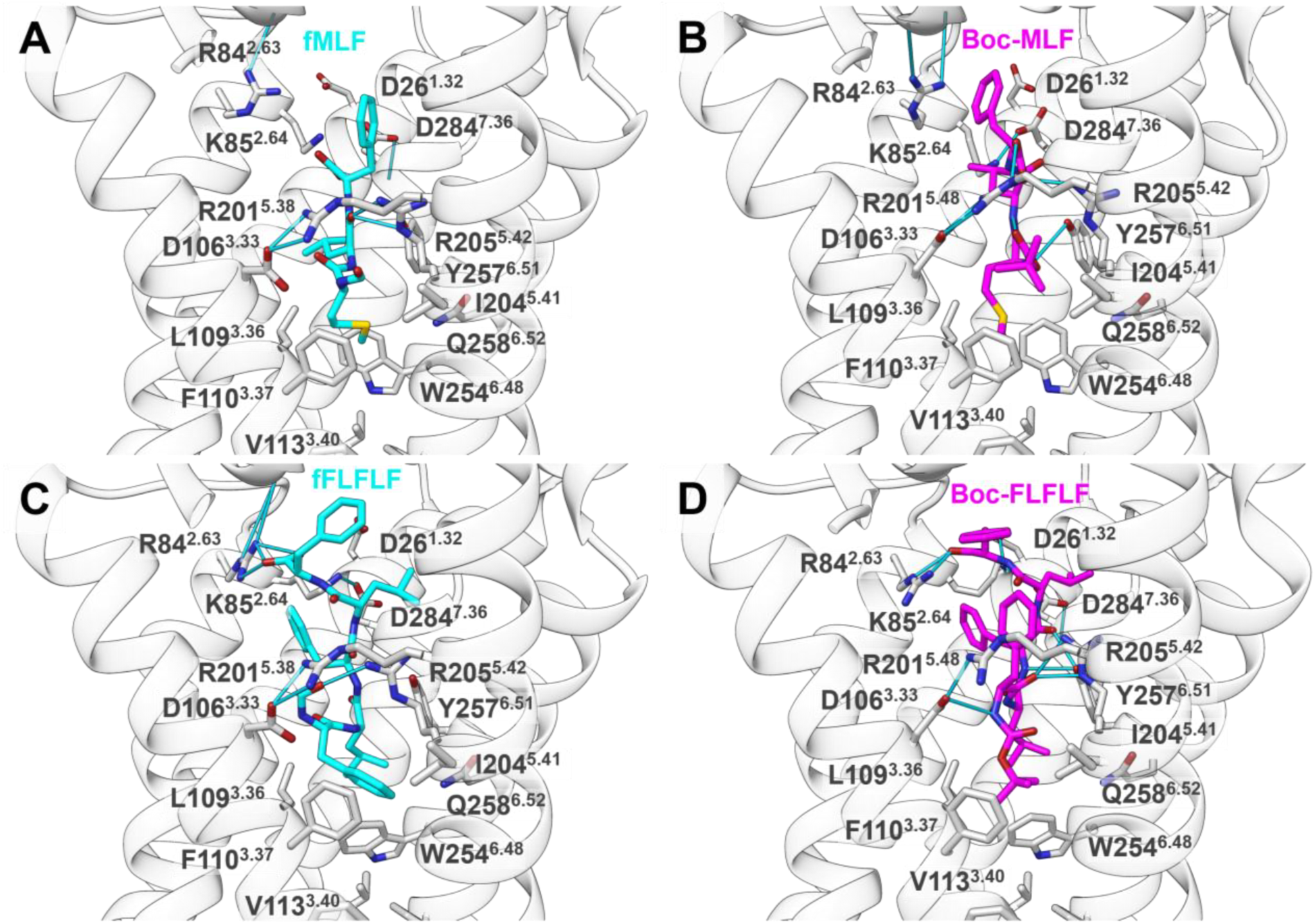
Binding mode of A) fMLF B) Boc-MLF C) fFLFLF and D) Boc-FLFLF.

In the course of the MD simulation, a dominant binding mode (25% of simulation frames) can be identified that preserves these key interactions. This includes a stable salt bridge between the negatively charged C-terminus of fMLF carboxylate with R84^2.63^ (∼70% of simulation frames; SI Figure S1A). In parallel, a cavity between TM3, TM4, and TM5 remained consistently accessible and is also observed in published FPR1 structures (TM3-TM5 hydrophobic cavity; Figure 2B). This observation led to the investigation of similar binding pockets in related GPCRs. In a study by Hedderich et al.^[26]^ allosteric sites were investigated by using a probe docking approach. A site that they identified named KS4 overlaps with the pocket that was found in FPR1 and can also be found in the cysteinyl leukotriene receptor (CysLT_1_R, PDB ID: 6RZ5^[27]^, SI Figure S5B) and the neurotensin receptor 1 (PDB ID: 6Z4S)^[28]^. CysLT_1_R (Uniprot ID Q9Y271) has a 27% sequence identity to FPR1 (Uniprot ID P21462). This cavity seems to occur in different GPCRs also with respect to inverse agonism.

We therefore hypothesised that this cavity accommodates the Boc group of the FPR1 antagonist Boc-MLF. Indeed, clustering of simulation frames identified a representative pose from the largest cluster (Figure 3B, SI Table S2). In this pose, the peptide backbone retains the agonist-like binding mode and maintains interactions with all three residues of the polar triad. The Boc group inserts into a cavity formed by the hydrophobic residues F110^3.37^, V113^3.40^ and I204^5.41^ and R205^5.42^ that is adjacent to the orthosteric site. Also, the Boc group causes a different orientation of the M1 and F1 side chain, which diminished engagement with the activation chamber. The insertion slightly expands the cavity (SI Figure S2) and is consistently observed in the dominant MD cluster.

Together, these results suggest that Boc-driven antagonism arises from occupation of a TM3-TM5 cavity that prevents the well-known conformational changes of TM3, TM5 and TM6 typically occurring upon activation of class A GPCRs.^[15,26]^ This mechanistic model is supported by docking and microsecond MD simulations and is consistent with available structural information on FPR1.

### Binding Mode Analysis of fFLFLF and Boc-FLFLF

Boc-FLFLF is another well-known antagonist. Based on the previous analysis, we hypothesised that fFLFLF would be the corresponding agonist. The subsequent computational analysis revealed that fFLFLF shows a similar binding mode as fMLF: The N-terminal formyl group engages the polar triad (∼30% of frames; SI Figure S3A), while the formylated phenylalanine side chain occupies the activation chamber via π-π interactions with F110^6.44^ and W254^6.48^ (Figure 3C) in a manner analogous to methionine. Additional hydrophobic contacts with residues L2–F5 of the peptide and receptor residues F33^1.39^, F74^2.53^, F81^2.60^, F102^3.29^, F178^4.52^, L78^2.57^, and V105^3.32^, support stable binding and activation. As with Boc-MLF, Boc-FLFLF preserves the peptide backbone interactions, while the Boc group inserts into a TM3-TM5 hydrophobic cavity (Figure 3D).

As an experimental validation, we employed a Homogeneous Time-Resolved Fluorescence (HTRF) cAMP assay^[18]^ to measure FPR1-mediated G_i_-dependent inhibition of *de novo* cAMP generation. Gi-dependent inhibition of de novo cAMP generation is a canonical FPR1 readout in HEK293 systems, with WKYMVm serving as a standard reference agonist. The HEK293-FPR1 stable line and HTRF cAMP-Gi assay provide a robust platform for quantifying FPR1 specific modulation. Consistent with its established functional profile, the formylated agonist fMLF inhibited cAMP accumulation with an E_max_ of 74 ± 3% and pEC_50_ of 8.2 ± 0.1, confirming full agonism. As predicted, fFLFLF is also an agonist with a pEC_50_ of 9.4 ± 0.1 and E_max_ of 77 ± 5%. In contrast, the corresponding Boc-coupled ligands Boc-MLF and Boc-FLFLF did not affect cAMP levels, consistent with the predicted binding modes. To directly test the antagonistic properties of the Boc-modified peptides, we used W-peptide as agonist. Within this experimental framework, Boc-MLF and Boc-FLFLF inhibited WKYMVm-induced Gi/cAMP signalling with pIC_50_ values of 5.8 ± 0.1 /7.3 ± 0.3 consistent with antagonism (Figure 4, Table 1).

**Table 1.** Dose-response curve parameters (logistic fit) of peptide derivatives.

| Ligand | $pEC_{50} \pm SEM$ | FPR1 response<br>$\pm SEM$ [%] | $pIC_{50} \pm SEM$ | Inhibition of<br>FPR1 Activity<br>$\pm SEM$ [%] |
| --- | --- | --- | --- | --- |
| fMLF | $8.2 \pm 0.1$ | $74 \pm 3$ | - | - |
| fFLFLF | $9.4 \pm 0.2$ | $77 \pm 5$ | - | - |
| fMLFYLA | $7.9 \pm 0.1$ | $70 \pm 4$ | - | - |
| Boc-MLF | ND | ND | $5.8 \pm 0.1$ | $71 \pm 5$ |
| Boc-FLFLF | ND | ND | $7.3 \pm 0.3$ | $55 \pm 8$ |
| Boc-MLFYLA | $5.6 \pm 0.2$ | $28 \pm 4$ | $7.8 \pm 0.2$ | $86 \pm 9$ |
| Fmoc-MLF | $6.6 \pm 0.5$ | $26 \pm 6$ | ND | ND |
| Fmoc-FLFLF | $4.7 \pm 1.6$ | > range* | ND | ND |
| Fmoc-MLFYLA | $6.6 \pm 0.4$ | $44 \pm 7$ | $7.9 \pm 0.2$ | $83 \pm 7$ |
| Trp-MLF | $6.4 \pm 0.09$ | $22 \pm 1$ | $8.2 \pm 1.2$ | $53 \pm 32$ |
| Trp-FLFLF | $5.6 \pm 0.4$ | $50? \pm 15$ | $6.5 \pm 0.6$ | $63 \pm 17$ |
| Trp-MLFYLA | $6.2 \pm 0.2$ | $71 \pm 7$ | $7.9 \pm 0.5$ | $45 \pm 11$ |

\*Estimated from model; exceeds experimental limits; ND = not detected

**Figure 4.**
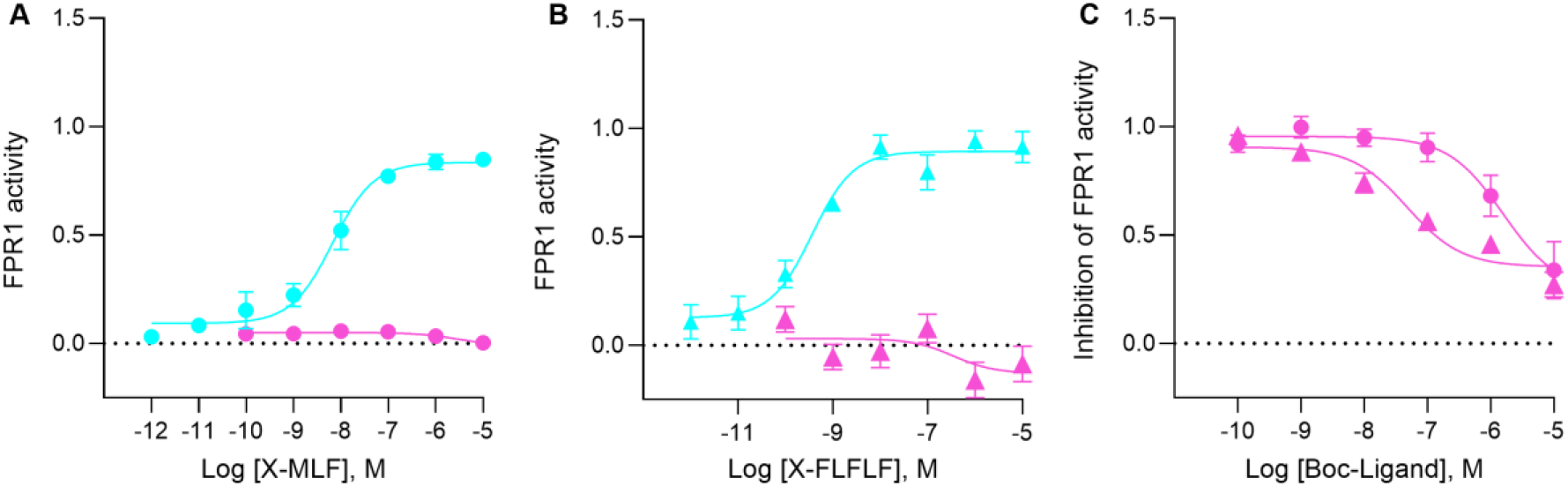
Pharmacological characterization of FPR1 ligands. A) Dose–response curves for fMLF (cyan) and Boc-MLF (magenta) showing agonist activity at FPR1. FPR1-expressing cells were treated with increasing ligand concentrations, and receptor activation was quantified as inhibition of forskolin-stimulated intracellular cAMP via Gαi signalling. Responses are reported as fraction of maximal inhibition of forskolin-induced cAMP. B) Dose– response curves for fFLFLF (cyan) and Boc-FLFLF (magenta) under the same conditions as in A). C) Antagonist activity of Boc-MLF (circles) and Boc-FLFLF (triangles), measured as inhibition of FPR1 activation by the agonist WKYMVm (W-peptide). Data are expressed as fraction of WKYMVm-induced response in the absence of antagonist. Data are mean ± SEM from five independent experiments (n = 5). Curves were fitted by nonlinear regression using a four-parameter logistic model with Hill slope constrained to 1.

### Analysis of structural requirements for antagonism

Based on the observations that Boc groups convert MLF/FLFLF peptides into antagonists by occupying a hydrophobic cavity between TM3 and TM5, we explored whether even larger hydrophobic modifications, such as Fmoc, can similarly confer antagonistic properties. MD simulations of Fmoc-MLF indicated a stable binding mode observed in 37% of frames (SI Table S2, Figure 5A). As with Boc, the Fmoc moiety inserts between TM3 and TM5, but extends further outwards of the receptor. The Fmoc group interacts with R201^5.38^ and R205^5.42^ in ∼20% and 35% of frames, respectively (SI Figure S1C).

**Figure 5.**
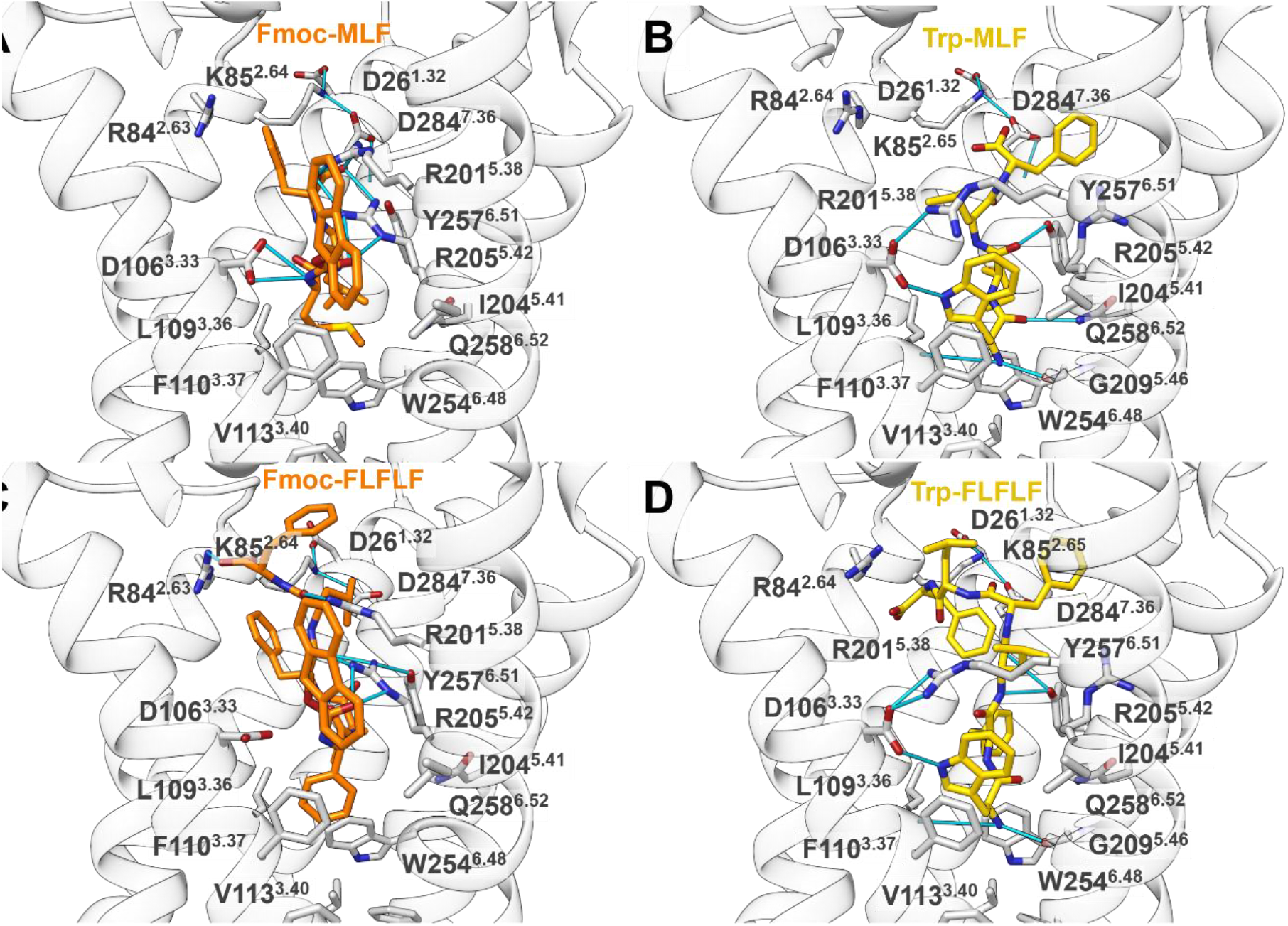
Binding mode of A) Fmoc-MLF B) Trp-MLF C) Fmoc-FLFLF and D) Trp-FLFLF.

Due to its flat aromatic structure, Fmoc disrupts the salt bridge between D106^3.33^ and R201^5.38^, while forming interactions with all three polar triad residues and Y257^6.51^. H-bonds with R84^2.63^ are observed only sporadically. Indeed, partial activation of FPR1 was detected with Fmoc-MLF (E_max_ of 26% and pEC_50_ 6.6) and Fmoc-MLF cannot block WKYMVm induced FPR1 activation, indicating no antagonistic effect in the tested concentrations (Table 1, Figure 6). In Fmoc-FLFLF, the Fmoc group inserts between TM3 and TM5, partially disrupting polar triad interactions, while the Fmoc carbamate forms contacts with D106^3.33^ in ∼35% of frames (Figure 5C, SI Figure S3C). The flat, bulky Fmoc aromatic system also extends toward TM4, allowing π-π stacking with F110^3.37^ and F114^3.41^. While Fmoc-FLFLF activated FPR1 with a pEC_50_ of 4.7, Fmoc-FLFLF could not antagonise FPR1 activation by W-peptide, similarly to Fmoc-MLF.

**Figure 6.**
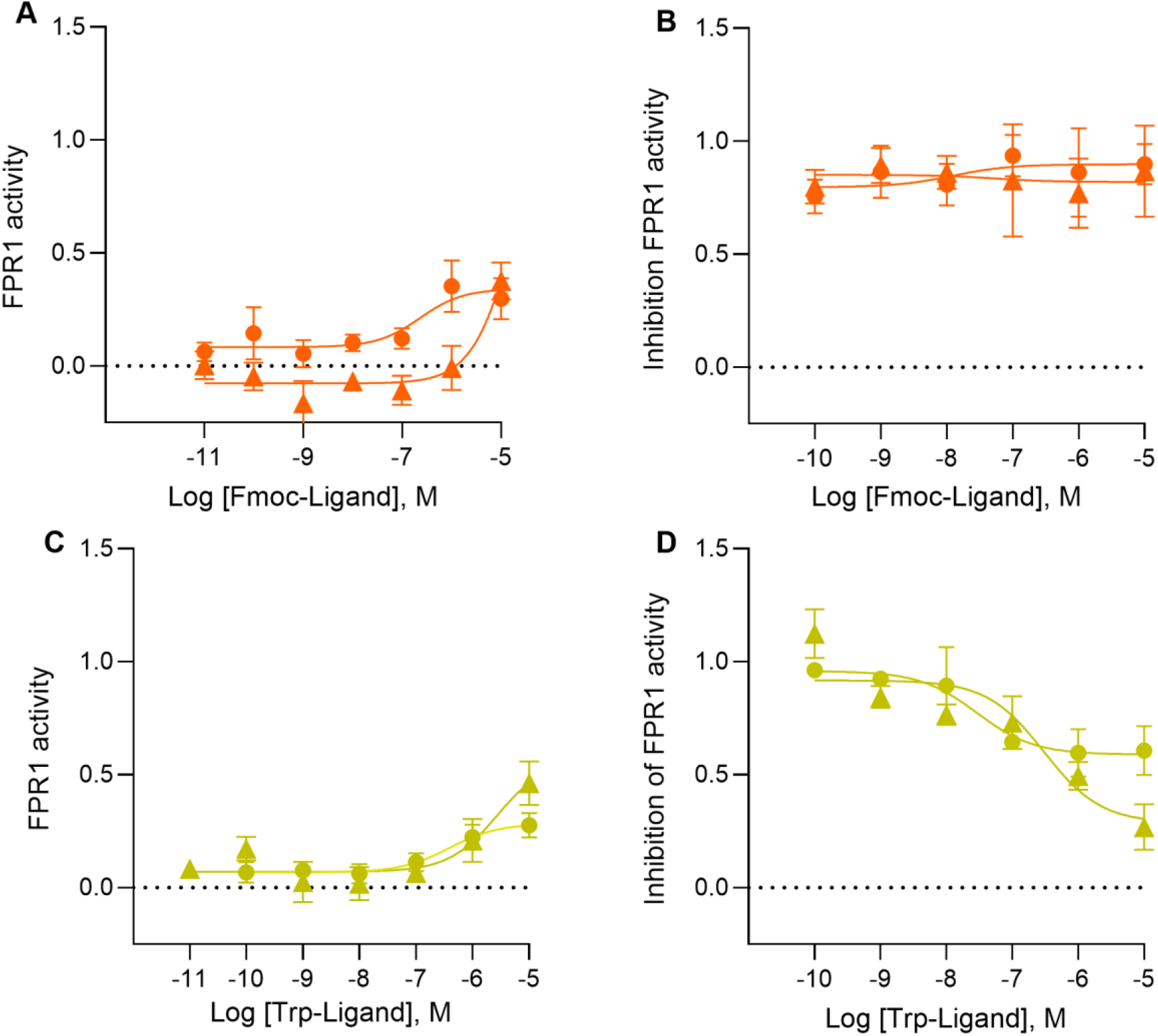
A) Dose–response curves for Fmoc-MLF (circles) and Fmoc-FLFLF (triangles) showing agonist activity at FPR1. FPR1-expressing cells were treated with increasing peptide concentrations, and receptor activation was quantified as inhibition of forskolin-stimulated intracellular cAMP via Gαi signalling. Responses are expressed as fraction of maximal inhibition of forskolin-induced cAMP. B) Antagonist activity of Fmoc-MLF and Fmoc-FLFLF, assessed by their ability to inhibit FPR1 activation induced by the agonist WKYMVm (W-peptide). Effects are expressed as fraction of WKYMVm-induced response in the absence of test ligand. C) Dose–response curves for Trp-MLF (circles) and Trp-FLFLF (triangles) evaluated under the same agonist assay conditions as in A). D) Antagonist activity of Trp-MLF and Trp-FLFLF determined using WKYMVm-induced FPR1 activation as described in B). Data are mean ± SEM from five independent experiments (n = 5). Curves were fitted by nonlinear regression using a four-parameter logistic model with Hill slope constrained to 1.

We next evaluated the effect of adding an N-terminal tryptophan as a bulky, hydrophobic moiety mimicking Boc to the MLF and FLFLF core. The most populated cluster (∼41% of frames) shows that the Trp side chain in Trp-MLF forms hydrogen bonds with the polar triad and additional interactions not observed for other MLF derivatives. Specifically, the Trp nitrogen acts as an H-bond donor to the D106^3.33^ backbone in ∼20% of frames, while the charged N-terminus interacts with the backbone oxygen of L109^3.36^ and G209^5.46^, maintaining an almost parallel orientation to TM3 and TM5 (Figure 5B). Additional H-bonds are formed with Q258^6.52^ and the backbone of R205^5.42^ in ∼25% of frames (SI Figure S1D). These results indicate that, even without the canonical N-formyl recognition motif, the Trp group can stabilise interactions with the polar triad while partially occupying the TM3-TM5 cavity. Functionally, Trp-MLF was also a weak FPR1 activator (pEC_50_ 6.4 ± 0.1, E_max_ 22 ± 1, Figure 6). The lack of strong agonism may be explained by the absence of ligand insertion into the activation chamber surrounding W254^6.48^, a highly conserved residue in Class A GPCRs described as the “rotameric switch” that transduces agonist binding to the G-protein binding site on the intracellular side of the receptor.^[29]^

However, Trp-MLF antagonised WKYMVm-induced receptor activation with a pIC_50_ of 8.2 ± 1.2 and an E_max_ of 53% (Figure 6, Table 1), consistent with the computational predictions. The Trp side chain nitrogen of Trp-FLFLF can form H-bonds with the polar triad. The positively charged N-terminus interacts with the backbone oxygen atoms of G209^5.46^ and L109^3.36^ (Figure 5D). These polar interactions position the Trp group almost parallel to TM3 and TM5, creating a unique binding mode that slightly increases the distance between these helices while still maintaining contacts with both, effectively keeping them in close proximity. Indeed, Trp-FLFLF activated FPR1 with a pEC_50_ of 5.6 ± 0.4 (E_max_ 50% ± 15), consistent with the computational observations that the Trp moiety stabilizes interactions with the polar triad while failing to engage the activation chamber fully. However, Trp-FLFLF is also able to partially inhibit WKYMVm activity with a pIC_50_ of 6.5 ± 0.6 and an E_max_ of around 63% ± 17 (Figure 6, Table 1).

### Extension of Structural Insights to fMLFYLA-based ligands

Building on insights from the MLF and FLFLF peptide series, we next investigated a third peptide series based on *Desulfotomaculum*-SP8 (fMLFYLA), a formylated hexapeptide from *Desulfotomaculum reducens*.^[30,31]^ The series included fMLFYLA, Boc-MLFYLA, Fmoc-MLFYLA, and Trp-MLFYLA, which were first analysed computationally. Overall, all four MLFYLA peptides adopt binding modes similar to the previously studied series, displaying comparable interaction patterns (Figure 7). Compared with the MLF and FLFLF peptides, MLFYLA has a longer sequence and additional H-bonding capabilities through the tyrosine side chain. MD simulations captured two conformations for this Y4 residue of fMLFYLA. In the first conformation, Y4 engages in π-π stacking with Y257^6.51^ and forms H-bonds with this residue in ∼35% of frames. In the second binding mode, Y4 forms an additional H-bond with D284^7.36^ (∼5% of frames), contributing to an extended polar network. This polar network is further stabilised by a salt bridge between K85^2.64^ and D284^7.36^, observed in ∼30% of simulation frames and consistently across all fMLFYLA simulations (Figure 7A, I Figure S3A).

**Figure 7.**
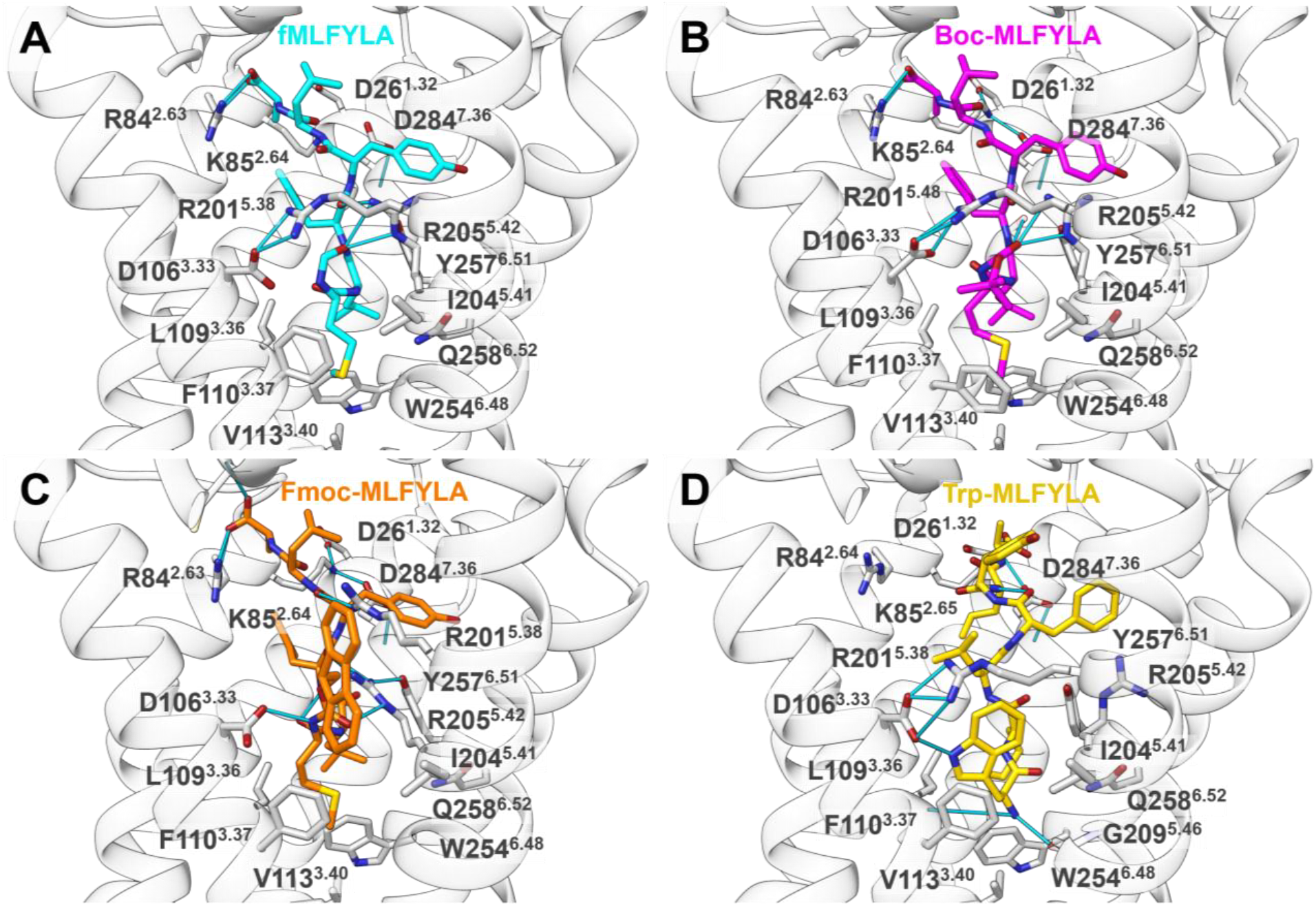
Binding modes of the MLFYLA (SP8) series: A) fMLFYLA B) Boc-MLFYLA C) Fmoc-MLFYLA and D) Trp-MLFYLA.

In the HTRF cAMP assay, fMLFYLA activated FPR1 with an E_max_ of 70% ± 4 and a pEC_50_ of 7.9 ± 0.1 (Table 1), in agreement with previous reports^[31]^ and confirming the predicted agonist activity.

All N-terminally modified peptides Boc-MLFYLA, Fmoc-MLFYLA, and Trp-MLFYLA partially activated FPR1 with efficacies of 28% ± 4, 44% ± 7, and 71% ± 7, respectively, and pEC_50_ values between 5.6-6.6 (Figure 8, Table1). In line with their nature as partial agonists, all three peptides also antagonised WKYMVm-induced FPR1 activation, with pIC_50_ values around 8 and E_max_ values ranging from 45-86%, supporting the predicted role of bulky N-terminal modifications in conferring antagonistic activity (Table 1).

**Figure 8.**
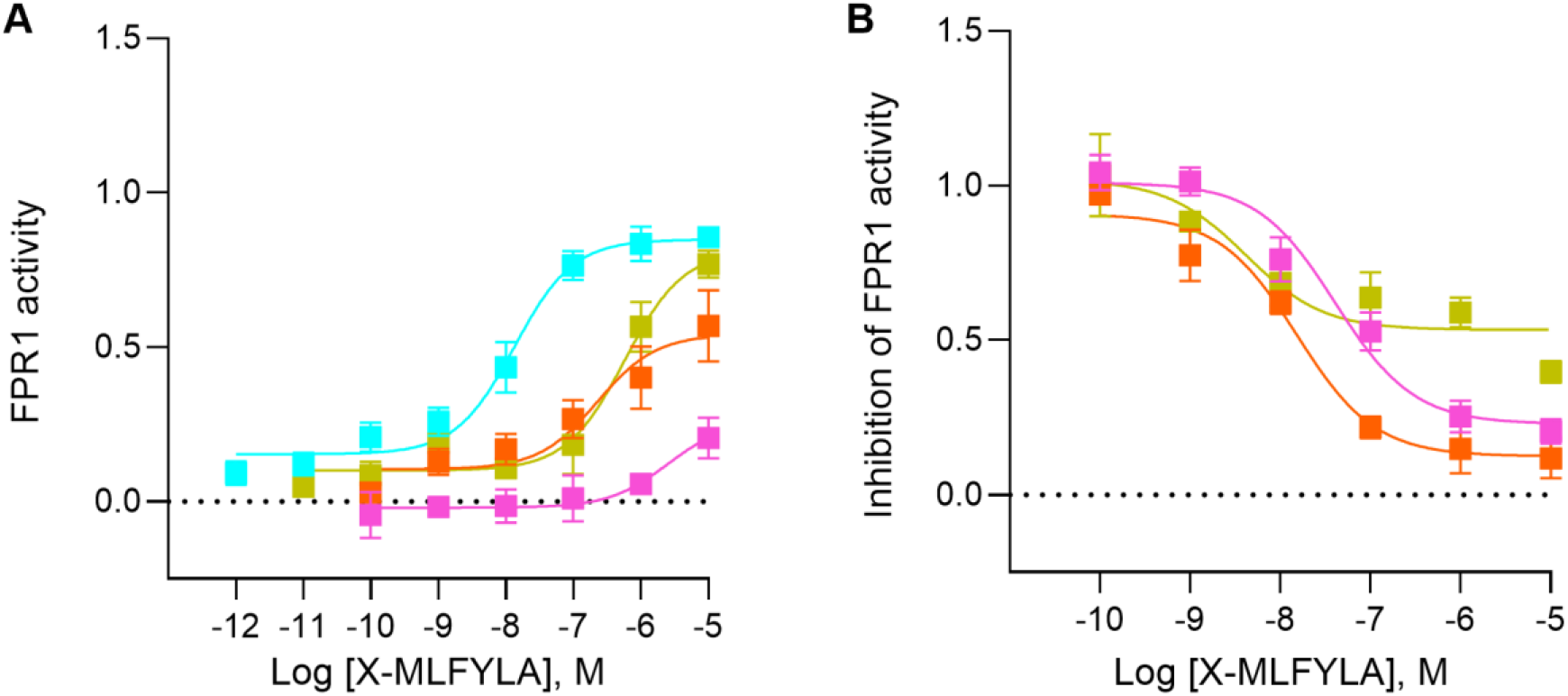
A) Dose-response curves of fMLFYLA (cyan circle), Boc-(pink square), Fmoc-(orange triangle) and Trp-MLFYLA (yellow diamonds. B) Boc-(pink squares), Fmoc-(orange triangles) and Trp-MLFYLA (yellow diamonds) determined by their ability to inhibit receptor activation induced by the FPR1 agonist W-peptide. Antagonist effects are expressed as fractions of WKYMVm-induced FPR1 in the absence of test ligand. For all panels, data represent mean ± SEM of five independent experiments (n = 5). Dose–response curves were fitted by nonlinear regression using a four-parameter logistic model with the Hill slope constrained to 1.

**Figure 9.**
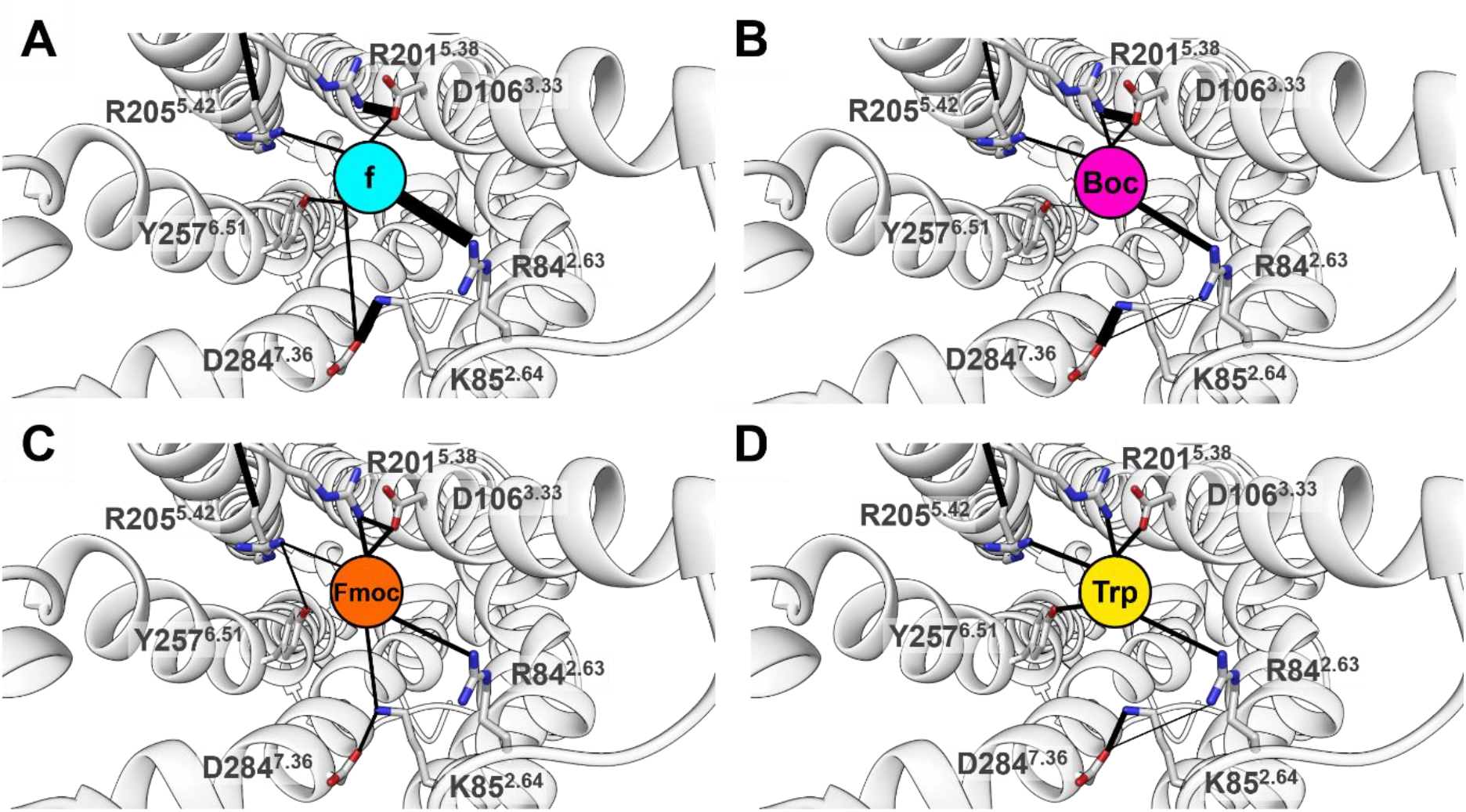
Summarised H-bond networks: A) Formylated (f) ligands, which are agonists. B) Ligands with a Boc group, which are mostly antagonist. C) Ligands featuring a Fmoc group or D) Trp group which mostly act as partial agonists. The black lines represent H-bonds. Line thickness reflects interaction frequency; thicker black lines represent more frequent interactions between residues.

## Discussion

A conserved interaction network involving R84^2.63^, the polar triad (D106^3.33^, R201^5.38^ and R205^5.42^) and Y257^6.51^ emerges as a key determinant of FPR1 activation, as it is engaged by all active ligands. All three share a formylated hydrophobic N-terminus (fM1 or fF1) that inserts into the activation chamber, with the formyl group anchoring binding via the polar triad, consistent with previous reports.

Previous docking studies suggested that the antagonist Boc-MLF binds high in the orthosteric site, with only limited interaction with the polar triad and no engagement of the activation chamber^[17]^. Our results indicate that Boc-driven antagonism arises from occupation of a TM3-TM5 cavity adjacent to the orthosteric site while preserving the canonical peptide binding mode. This mechanistic model is supported by docking, microsecond MD simulations, and pharmacological data, and is consistent with available FPR1 structures. Insertion of hydrophobic groups into the TM3-TM5 hydrophobic cavity was already shown for FPR2, where an aliphatic side chain of the activator Quin-C1 is inserted into this pocket (SI Figure S5C).^[32]^ Accordingly, the antagonistic properties are consistent with insertion of the Boc group into a hydrophobic cavity between TM3 and TM5, pending direct structural validation.

Comparison of the two short backbones (MLF and FLFLF) and one longer hexapeptide (MLFYLA) shows consistent placement of the Boc group within the TM3-TM5 cavity, accompanied by clear antagonism of Boc-modified ligands (Table 1). In this broader context, hydrophobic N-terminal modifications across MLF, FLFLF, and MLFYLA generally reduce receptor activation consistent with occupation of a pocket adjacent to the orthosteric site, but with variable functional outcomes ranging from full antagonism to partial agonism. Thus, the Boc-mediated agonist-to-antagonist conversion appears transferable across peptide sequences, as demonstrated for the tri- and pentapeptide series. Together, these findings support a general mechanism and indicate sequence- and length-dependent effects. Systematic exploration of broader sequence diversity will be required to define the limits of this mechanism.

We therefore also investigated the effect of the addition of the bigger group Fmoc to the three peptide series. The computational studies suggest that Fmoc insertion into the TM3-TM5 hydrophobic pocket may weaken the R201^5.38^-D106^3.33^ salt-bridge. The breakage of such an important interaction might explain the low pEC_50_ for Fmoc-MLF and Fmoc-FLFLF. As the activity of these peptides is very low, they are probably not able to replace W-peptide in the tested concentrations. Fmoc-MLFYLA on the other hand has one more residue and Y4 shows additional π-π stacking interactions with aromatic receptor residues. This might explain its higher activity and ability to replace W-peptide. Fmoc-MLFYLA has a very high antagonist efficacy but also partially activates the receptor which classifies it as a partial agonist.

The Trp derivatives have a free N-terminus which could impede the binding considering the high abundance of basic residues like R84^2.63^, K85^2.64^, R201^5.38^ and R205^5.42^ in the orthosteric site. However, the computational results suggested that for these ligands the nitrogen of the Trp side chain forms interactions with D106^3.33^, shielding the ligand N-terminus from R201^5.38^ and pushing R205^5.42^ away from the centre of the orthosteric site (Figure 5B,D and 7D). The *in vitro* results are consistent with this hypothesis. Moreover, all three ligands are able to replace W-peptide and can therefore be classified as partial agonists (Table 1).

In conclusion, our combined computational and *in vitro* approach was able to show that N-terminal modifications of peptides binding to FPR1 influence their activity. In general, hydrophobic groups interfere with receptor activation likely through insertion in a pocket adjacent to the orthosteric site that hinders full engagement of the activation chamber.

## Methods

### FPR1 homology modelling

The crystal structure of the human formyl peptide receptor 2 (FPR2) in the active state in complex with WKYMVm (PDB ID 6LW5^[16]^) was used as a template for homology modelling of FPR1, given the high sequence similarity between FPR1 and FPR2 of 69%.^[14]^ Structure preparation was performed using Molecular Operating Environment (MOE) version 2019.^[33]^ First, the FPR2 structure was prepared by removing the thermostabilising apocytochrome introduced at the N-terminus and adding the missing N-terminal residues M1 and E2. The missing C-terminal residues E347-M350 were also added. The introduced mutation S211^5.48^ was reverted to the native leucine residue. Bound cholesterol and water molecules were removed. Protonation states were assigned using the “3D Protonate” module at pH 7.4 in MOE. The structure was then energy-minimized using the AMBER10:EHT force-field with positional restraints applied to preserve structural integrity.

The sequences of FPR1 and the prepared FPR2 template were aligned using MOE, and 100 homology models of FPR1 were generated. The top-ranking model based on the Generalized Born/Volume Integral (GB/VI) scoring function was selected for further refinement. Ten intermediate models were subsequently generated, and the highest scoring model was selected as the final refined structure. The quality of the final apo-FPR1 model was assessed based by visual inspection and evaluation of backbone geometry with no inconsistencies in torsion angles observed.

### Molecular docking

The binding site of WKYMVm in FPR2 is defined by the crystal structure supported by mutagenesis data.^[16]^ To validate the docking protocol, WKYMVm was first docked into the FPR1 homology model using Gold version 5.7.1.^[34,35]^ Docking poses of WKYMVm were ranked using Goldscore and rescored with ChemPLP. An automatic search efficiency of 200% was applied, and the options “flip pyramidal N”, “flip amide bond” and “flip ring corners” were enabled. The binding site in the FPR1 homology model was defined based on the position of the C-terminal residues of WKYMVm in FPR2 crystal structure, using a 5 Å radius sphere. The docking protocol was considered validated as it reproduced a binding mode consistent with the experimental FPR2 structure. Subsequently, all other ligands were docked using the same protocol. The best docking poses were based on: i) a plausible orientation within the binding site and ii) a low root-mean-square deviation (RMSD) relative to the backbone conformation of WKYMVm in the FPR2-bound crystal structure.

### Molecular dynamics simulation

Molecular dynamics (MD) simulations of FPR1 were performed using the YASARA package, following standard protocols.^[36,37]^ A simulation cell was constructed with 15 Å padding in the XZ-plane (membrane) and 10 Å along the Y-axis (solvent). A built-in script was used to identify exposed hydrophobic residues and orient the receptor in the membrane accordingly. The membrane was composed of 1-palmitoyl-2-oleoyl-sn-glycero-3-phosphocholine (POPC) and, the final orientation was assessed by visual inspection. Simulations were carried out at physiological ion strength (0.9% NaCl). The system comprised 183 lipids, 14,191 water molecules, 84 chloride, and 68 sodium ions. The temperature was set to 298 K and the pressure to 1 bar. Amber ff14SB^[38]^ was used for the protein, Lipid14^[39]^ for the lipids, and TIP3P^[40]^ for water molecules. Ligand parameters were generated with AM1-BCC^[41]^ and GAFF^[42]^. A 2*1.25 fs time step was used and the SHAKE algorithm^[43]^ was applied. Long-range Coulomb interactions were calculated using the Particle Mesh Ewald algorithm.^[44]^ and periodic boundary conditions were applied. The system was equilibrated for 250 ps and coordinates were saved every 100 ps.

The protocol described above was used to run MD simulations of FPR1 in complex with the investigated ligands (SI Table S3). Production simulations were run for minimal total simulation time of 1 µs and at least two independent simulations were performed. RMSD of ligands and receptor in the MD simulations were assessed based on the YASARA output. All trajectories were visualised using VMD 1.8.8^[45]^, and H-bond analysis was performed with the corresponding VMD plugin. H-bond analysis focused on the interactions between the respective ligand and key residues (D26^1.32^, R84^2.63^, R85^2.64^, D106^3.33^, R201^5.38^, R205^5.42^, Y257^6.51^ and D284^7.36^). H-bonds with an occurrence below 1% were excluded from further analysis. The plots and H-bond networks were created with in-house python scripts. The trajectories were clustered using cpptraj^[46]^ with the k-means algorithm. Ten representative structures were generated for each ligand-receptor complex. Representative clusters and populations are provided in the Supporting Information (SI Table S2). Representative structures were visualised with UCSF Chimera.^[47]^

### Peptide Ligands

Peptide agonists and antagonists used in this study were obtained from commercial suppliers or synthesised as specified below. All peptides were dissolved in the appropriate solvent, aliquoted, and stored according to the manufacturers’ recommendations to preserve stability and activity.

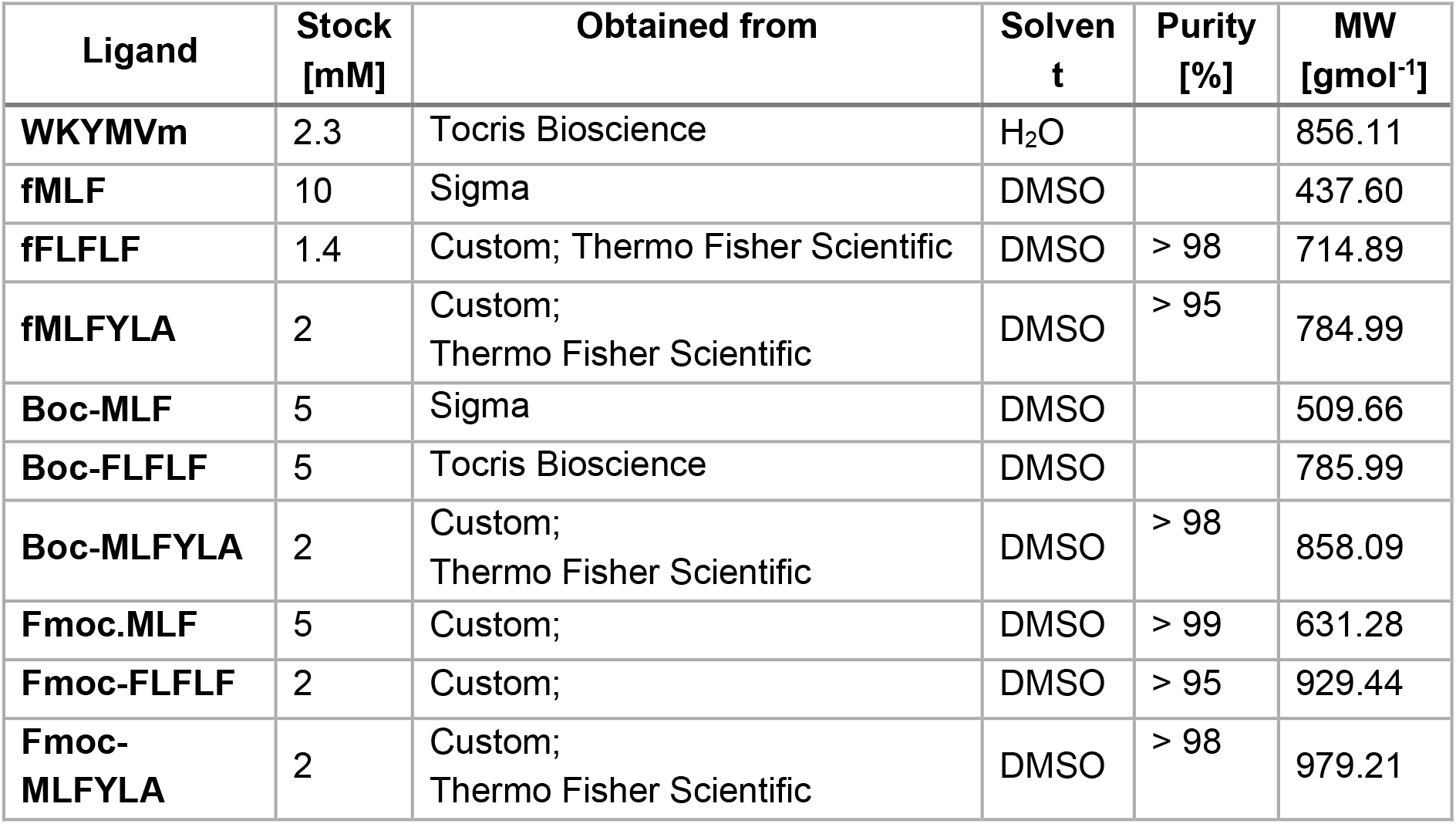
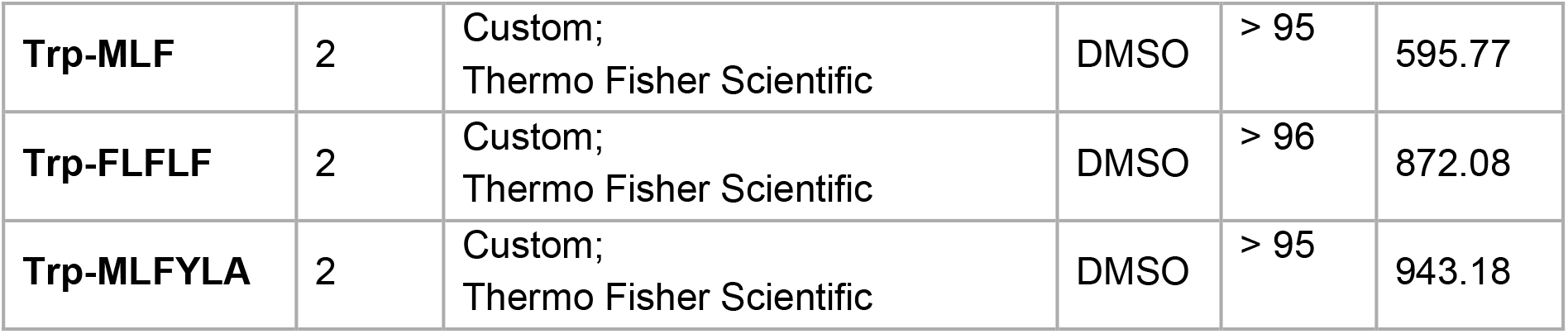

### Cell Culture

Human embryonic kidney (HEK293) cells, including the HEK293-FPR1 stable cell line generated previously described in Gröper et al.^[18]^ and Pajonczyk et al.^[19]^, were cultured in Dulbecco’s Modified Eagle’s Medium (DMEM; PAN-Biotech, Germany) supplemented with 10% standardized fetal bovine serum (FBS Advanced; Capricorn Scientific, Germany), 100 U/mL penicillin, 0.1 mg/mL streptomycin (GE Healthcare, USA), 1% L-glutamine, and 1% non-essential amino acids (NEAA; Sigma-Aldrich). Cells were maintained at 37°C in a humidified atmosphere containing 7% CO_2_.

### HTRF-based quantification of cAMP levels

Cells were seeded in 96-well plates, cellular *de novo* cAMP generation was activated through forskolin (FSK; Sigma-Aldrich), and cAMP levels were quantified using a Homogeneous Time-Resolved Fluorescence (HTRF) competitive immunoassay (cAMP-G_i_ Kit, Cisbio) as previously described. To assess antagonist activity, cells were pre-incubated with the respective ligands for 15 min prior to stimulation with W-peptide. Samples were then transferred to a 384-well plate and HTRF detection reagents were added. Fluorescence signals were measured using a CLARIOstar reader (BMG Labtech) with the following settings: 200 flashes per well, integration start 60 µs, integration time 400 µs, and settling time 100 µs. Signals were calculated as the ratio of 10,000 × (acceptor signal/donor signal) and normalized to the maximal system response, defined either by forskolin-induced cAMP accumulation (for verification of agonistic activity at FPR1) or by W-peptide-induced cAMP reduction (for assessment of antagonistic activity).

### Curve Fitting and Statistical Analysis

Data were normalized and expressed as fractions of the maximal system response. Curve fitting and analysis, including EC_50_, IC_50_ and E_max_ values, were performed using GraphPad Prism version 8.4.3 with a four-parameter sigmoidal model and a fixed Hill slope of 1. Experiments were normalized individually and dose-response curves were generated from at least five independent experiments. Data are presented as mean values ± standard error of the mean (SEM).

## Acknowledgments

This work was supported by grants from the German Research Foundation [DFG, SFB1009/A06] and the Interdisciplinary Center for Clinical Research of the Münster Medical Faculty [IZKF, Re2/014/26] to UR. J.M. and S.M. were funded by the German Research Foundation (GRK 2515 Chembion). O.K. was funded by a Heisenberg-Professorship (DFG; KO 4689/5-1 & KO 4689/5-2).

## Conflicts of Interest

O.K. is a scientific advisor at Nuvisan ICB GmbH and Prosion GmbH. The other authors declare no conflicts of interest.

## Data Availability Statement

The data that support the findings of this study are available on request.

## Author Contributions

O.K., U.R. and C.A.R. conceived the project, designed the experiments, and supervised the project. D.P. performed the pharmacological experiments. S.M. performed the computational experiments. S.M., D.P., J.M. C.A.R., U.R. and O.K. analyzed the data. T.B. assisted with peptide synthesis, purification, and characterization. All authors wrote and revised the manuscript. All authors have approved the final version of the manuscript.

