## Supporting Information for "Modulation of the agonist and antagonist activity of peptidic FPR1 ligands through N-terminal modifications: A structural and functional analysis"

Table S1: Overview about the available cryo-EM structures of FPR1 and the RMSD to the homology model generated in this study. The RMSD was calculated over the backbone atoms of 296 matching residue pairs of the 6LW5-based homology model and the respective experimental structure.

| PDB ID | Ligand | Resolution [Å] | Reference | RMSD to Homology Model [Å] |
|---|---|---|---|---|
| 7EUO | fMLF | 2.9 | Chen et al. 2022[1] | 1.92 |
| 7T6T | fMLFII | 3.2 | Zhuang et al. 2022[2] | 1.86 |
| 7VFX | fMIFL | 2.8 | Chen et al. 2022[1] | 1.93 |
| 7WVU | fMLF | 3.3 | Zhu et al. 2022[3] | 2.01 |

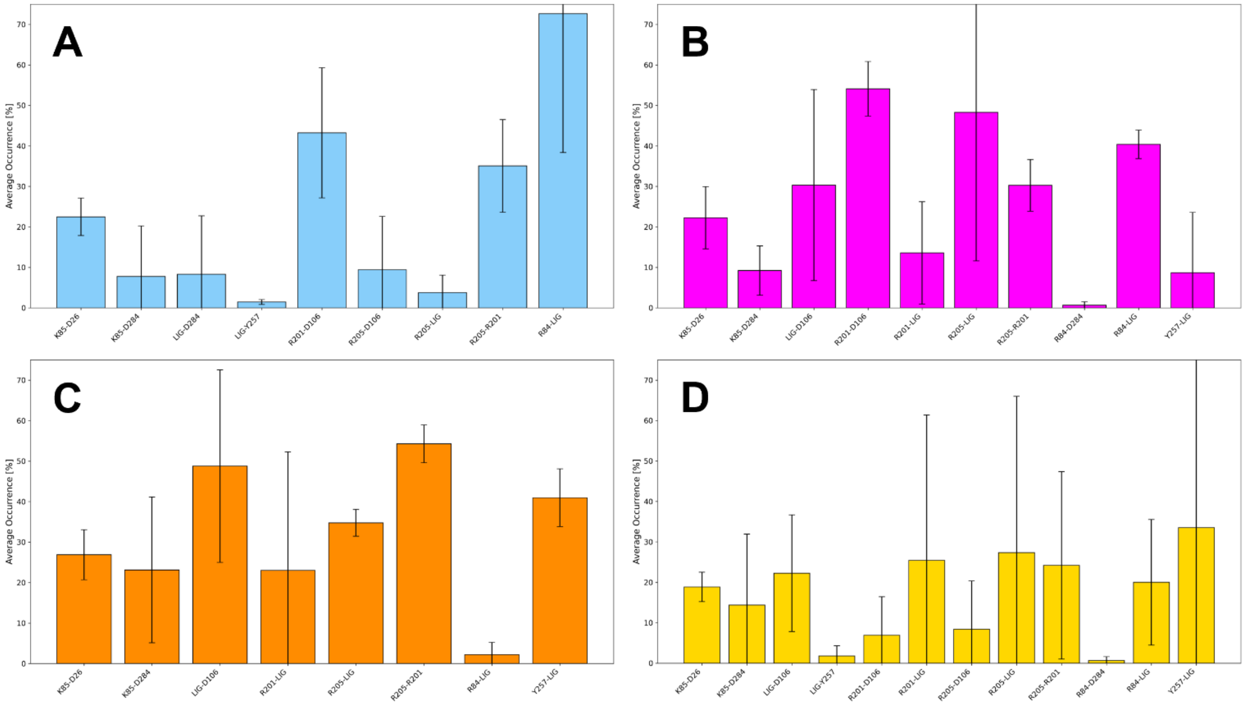

Figure S1: Hydrogen bonding interactions of MLF series (A) fMLF (B) Boc-MLF (C) Fmoc-MLF (D) Trp-MLF.

Table S2: Overview of clusters based on cpptraj[4] clustering of MD simulation frames of FPR1 in complex with the respective ligand.

| Peptide | Clusters | Frames in Most Populated Cluster | Frames in Most Populated Cluster |
|---|---|---|---|
| fMLF | 10 | 3721 | 25% |
| Boc-MLF | 10 | 2930 | 20% |
| Fmoc-MLF | 10 | 3705 | 37% |
| Trp-MLF | 10 | 4120 | 41% |
| fFLFLF | 10 | 4434 | 30% |
| Boc-FLFLF | 10 | 4439 | 44% |
| Fmoc-FLFLF | 10 | 4005 | 40% |
| Trp-FLFLF | 10 | 3267 | 22% |
| fMLFYLA | 10 | 4926 | 49% |
| Boc-MLFYLA | 10 | 5353 | 36% |
| Fmoc-MLFYLA | 10 | 4482 | 30% |
| Trp-MLFYLA | 10 | 2722 | 18% |

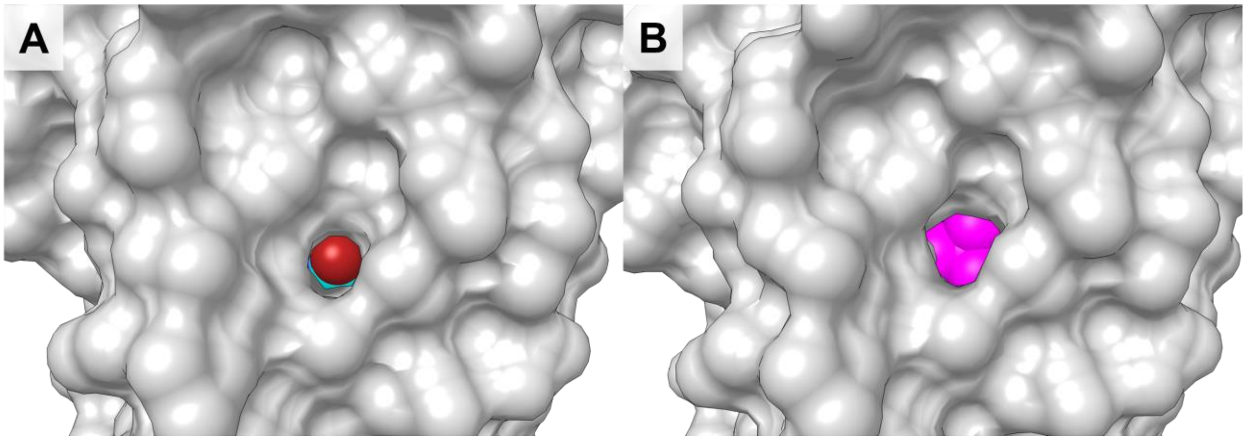

Figure S2: Hydrophobic cavity TM3-TM5 comparison between (A) fMLF and (B) Boc-MLF.

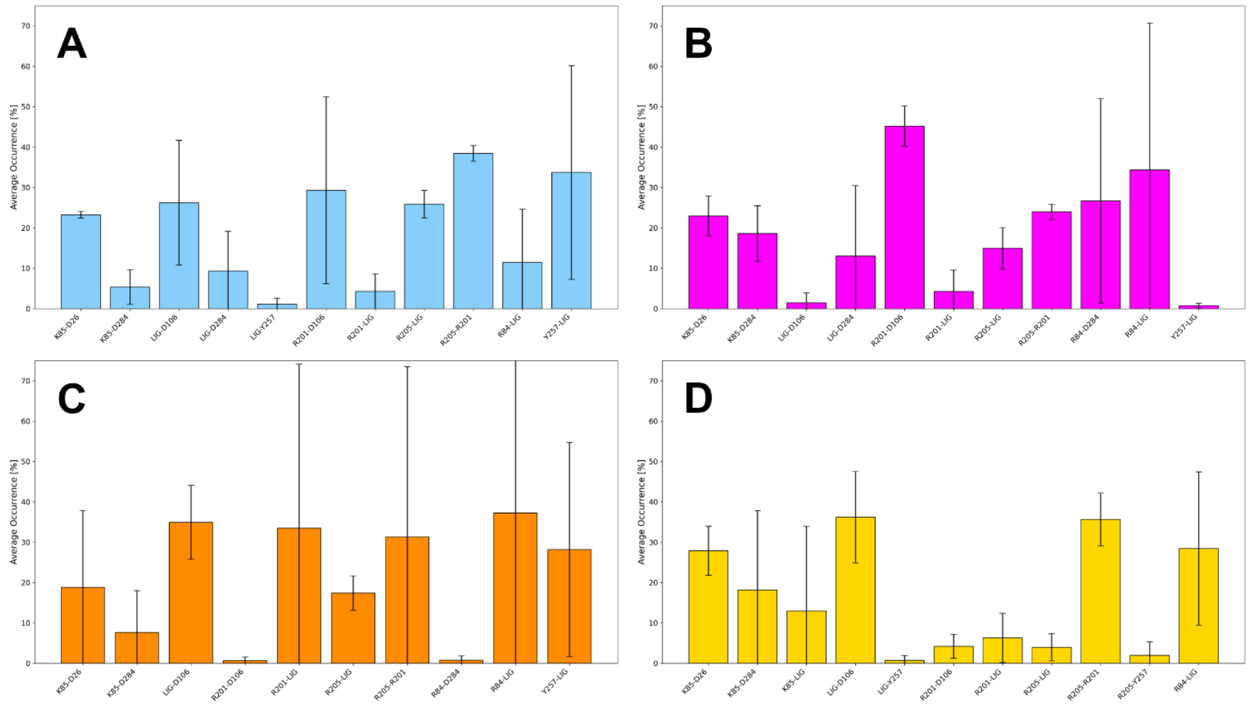

Figure S3: Hydrogen bonding interactions of FLFLF series (A) fFLFLF (B) Boc-FLFLF (C) Fmoc-FLFLF (D) Trp-FLFLF.

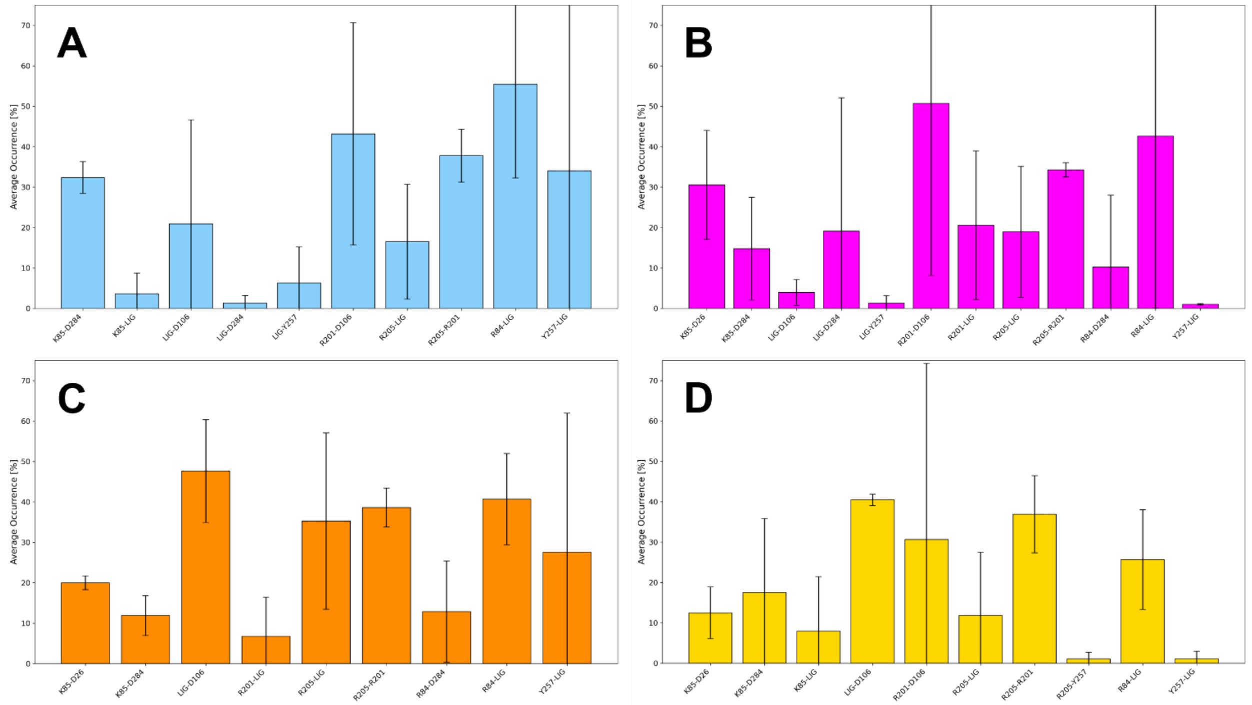

Figure S4: Hydrogen bonding interactions of MLFYLA series (A) fMLFYLA (B) Boc-MLFYLA (C) Fmoc-MLFYLA (D) Trp-MLFYLA

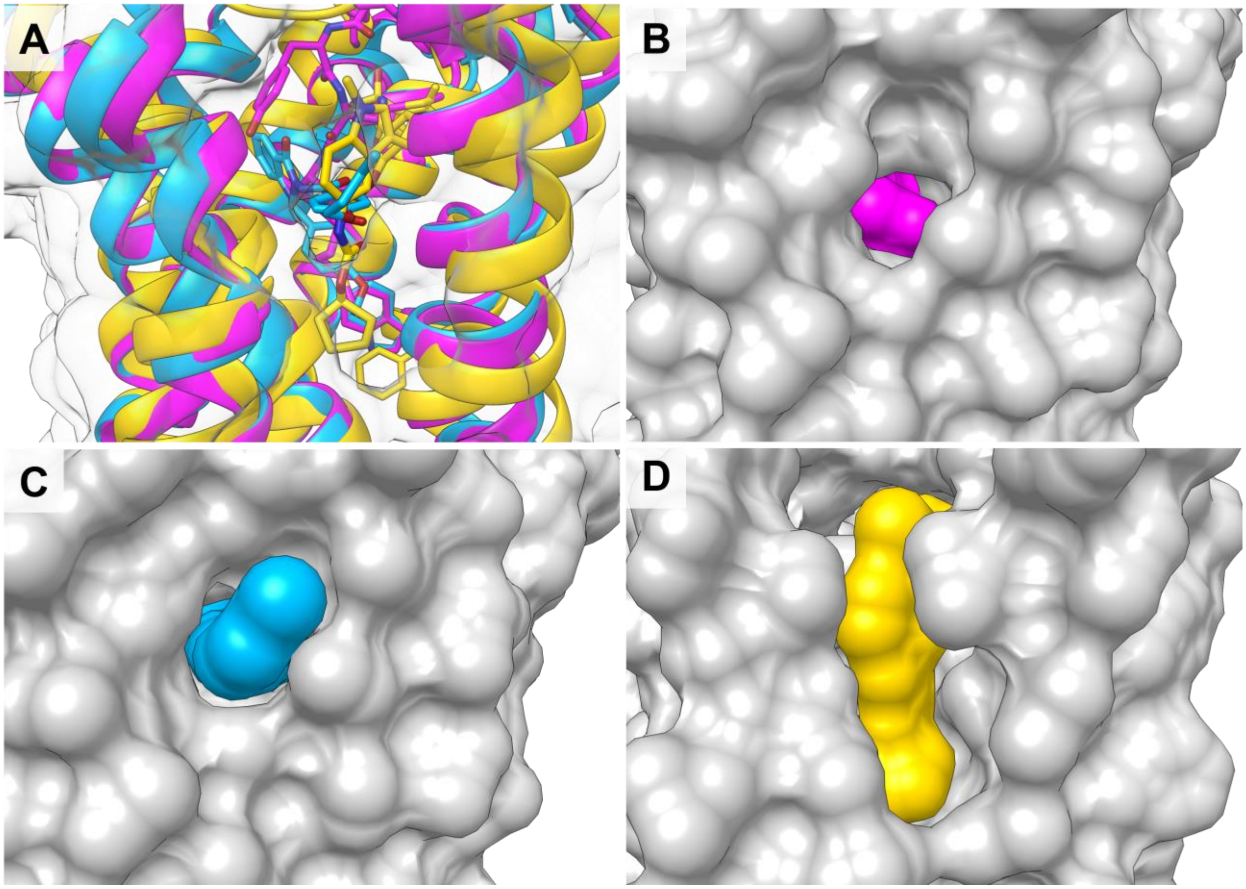

Figure S5: (A) Structural overlay of $CysLT_1R$ bound to the antagonist pranlukast (yellow, PDB ID: 6RZ5[5]), WKYMVm-$NH_2$-bound FPR2 (pink, PDB ID: 6LW5[6]) and Quin-C1-bound FPR2 (blue, PDB ID: 8ZBW[7]). (B) Solid surface depiction of FPR2 (grey) and the hydrophobic cavity imagined to be exploited by modifying known FPR1-ligands. WKYMVm-$NH_2$ is coloured pink. (C) Same pocket as in (B) but for Quin-C1-bound (blue) FPR2 (grey). (D) Similar pocket as in (B) but for pranlukast-bound (yellow) $CysLT_1R$ (grey).

Table S3: Overview of molecular dynamics (MD) simulations of FPR1 in complex with the respective ligand.

| Peptide | Analysed simulations | Total Simulation Length [µs] | Number of Atoms |
|---|---|---|---|
| fMLF | 3 | 1.5 | 56,982 |
| Boc-MLF | 3 | 1.5 | 55,966 |
| Fmoc-MLF | 2 | 1.0 | 59,717 |
| Trp-MLF | 2 | 1.0 | 57,126 |
| fFLFLF | 3 | 1.5 | 63,403 |
| Boc-FLFLF | 2 | 1.0 | 57,035 |
| Fmoc-FLFLF | 2 | 1.0 | 57,279 |
| Trp-FLFLF | 3 | 1.5 | 57,033 |
| fMLFYLA | 2 | 1.0 | 57,121 |
| Boc-MLFYLA | 3 | 1.5 | 57,104 |
| Fmoc-MLFYLA | 3 | 1.5 | 60,236 |
| Trp-MLFYLA | 3 | 1.5 | 57,119 |

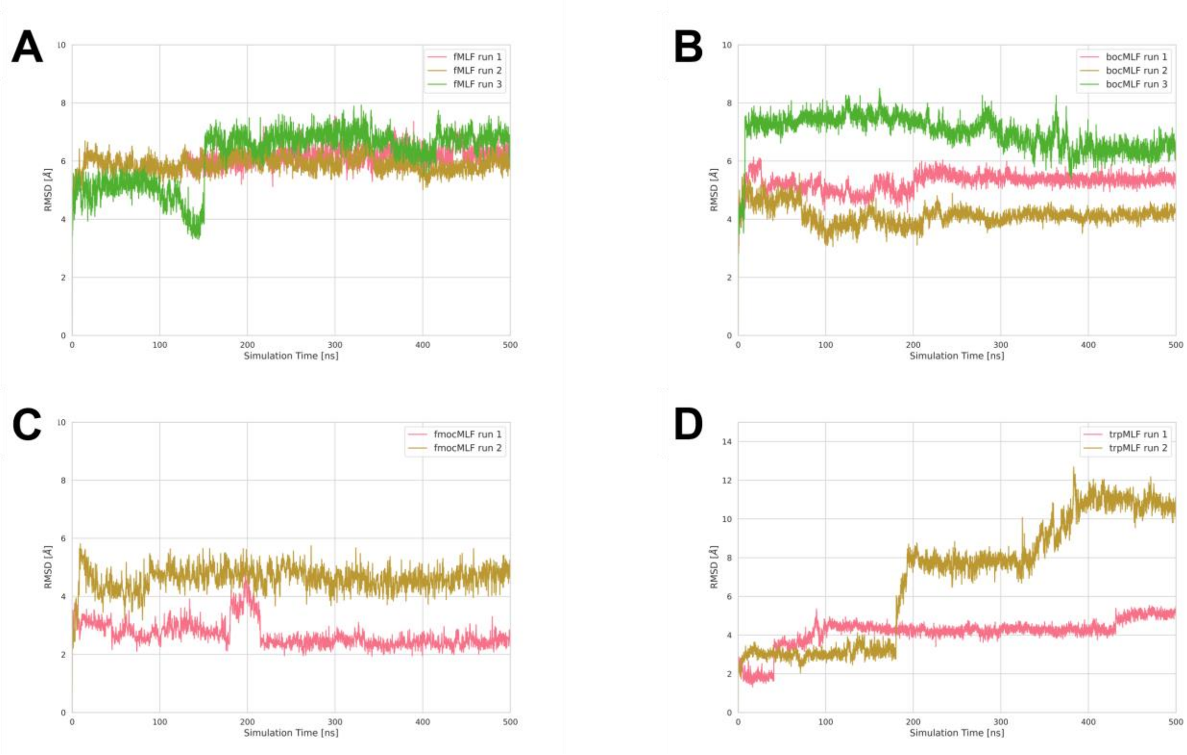

Figure S6: RMSD of all heavy atoms of the ligands of the MLF series. (A) fMLF (B) Boc-MLF (C) Fmoc-MLF (D) Trp-MLF

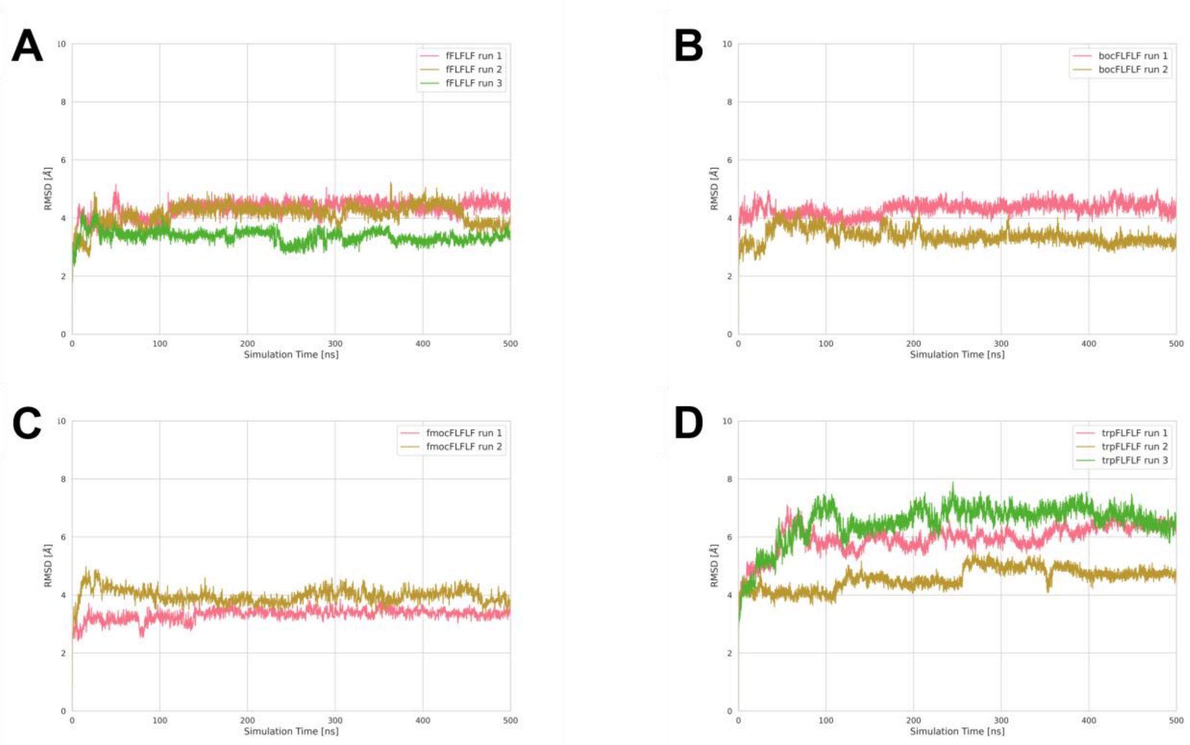

Figure S7: RMSD of all heavy atoms of the ligands of the FLFLF series. (A) fFLFLF (B) Boc-FLFLF (C) Fmoc-FLFLF (D) Trp-FLFLF

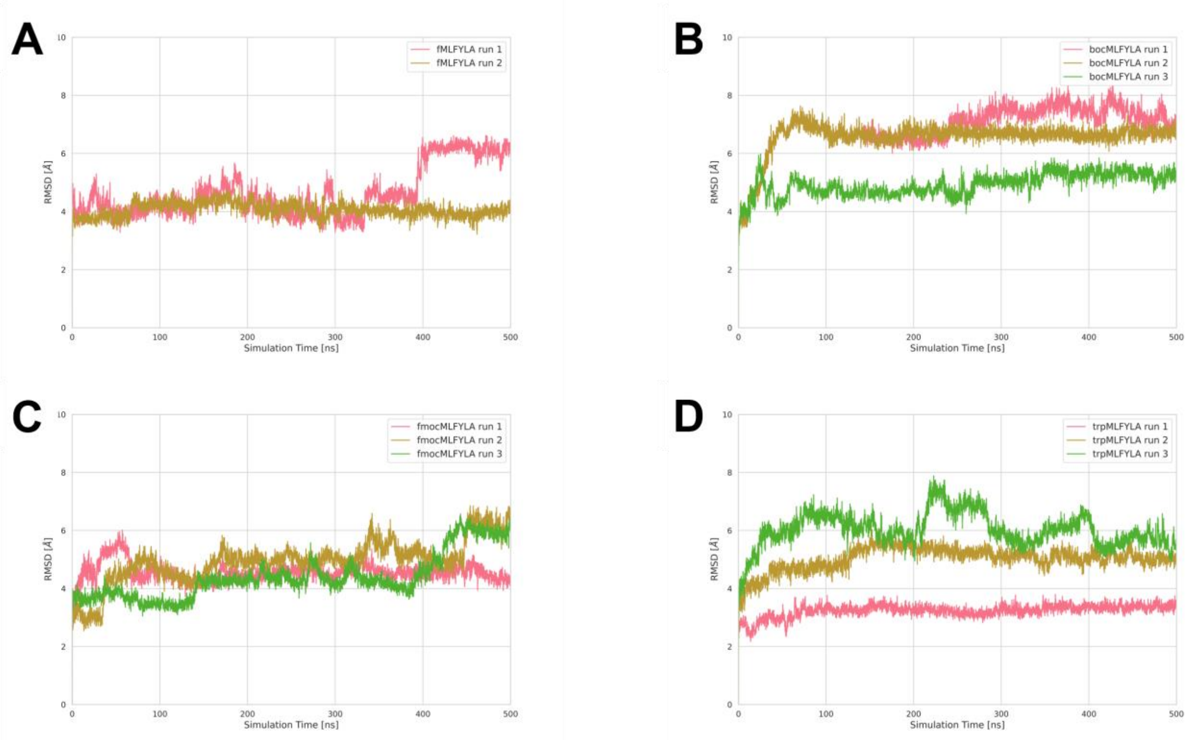

Figure S8: RMSD of all heavy atoms of the ligands of the MLFYLA series. (**A**) fMLFYLA (**B**) Boc-MLFYLA (**C**) Fmoc-MLFYLA (**D**) Trp-MLFYLA

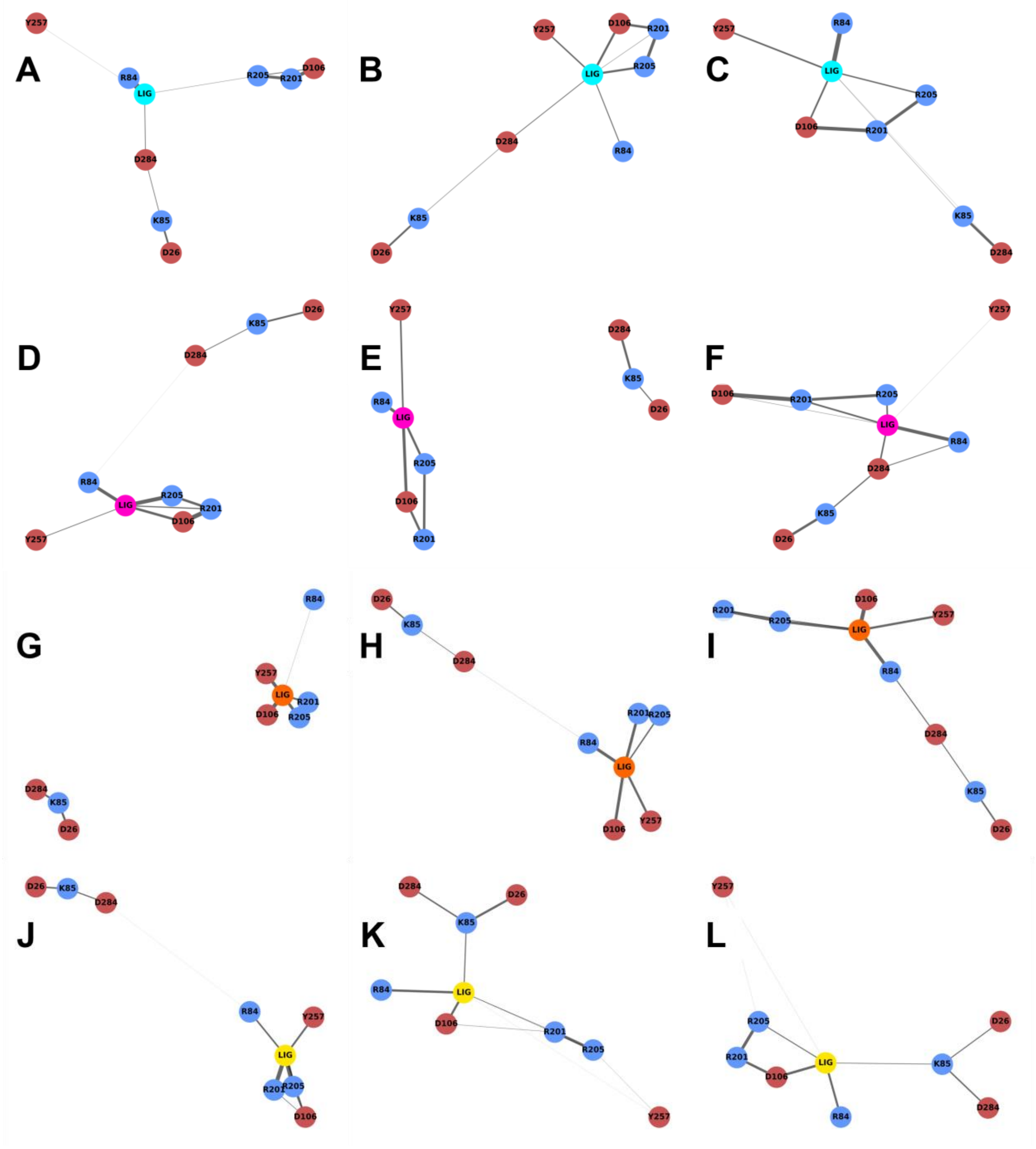

Figure S9: Hydrogen bond networks of (A) fMLF (B) fFLFLF (C) fMLFYLA (D) Boc-MLF (E) Boc-FLFLF (F) Boc-MLFYLA (G) Fmoc-MLF (H) Fmoc-FLFLF (I) Fmoc-MLFYLA (J) Trp-MLF (K) Trp-FLFLF (L) Trp-MLFYLA

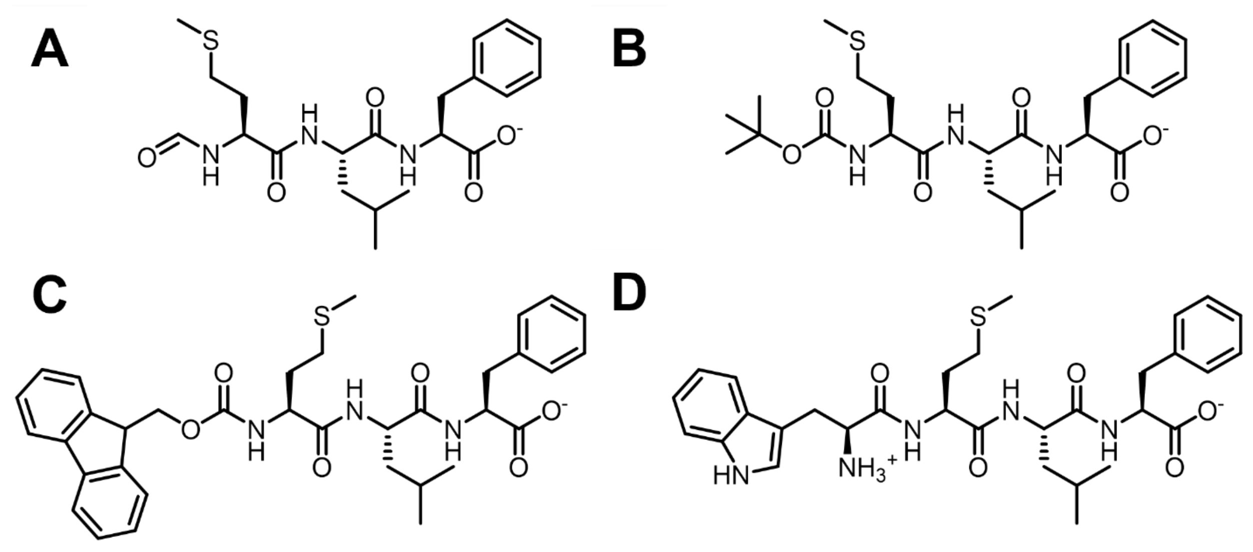

Figure S10: Structures of MLF series (A) fMLF (B) Boc-MLF (C) Fmoc-MLF (D) Trp-MLF.

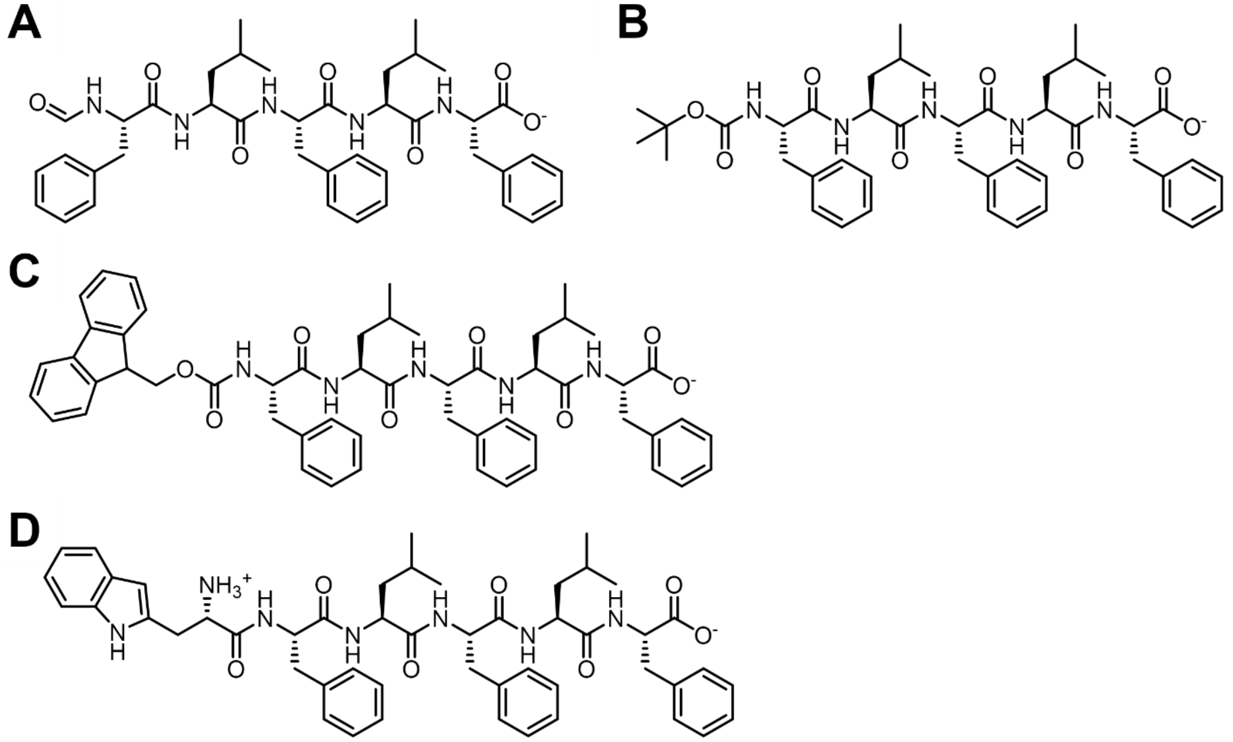

Figure S11: Structures of FLFLF series (A) fFLFLF (B) Boc-FLFLF (C) Fmoc-FLFLF (D) Trp-FLFLF.

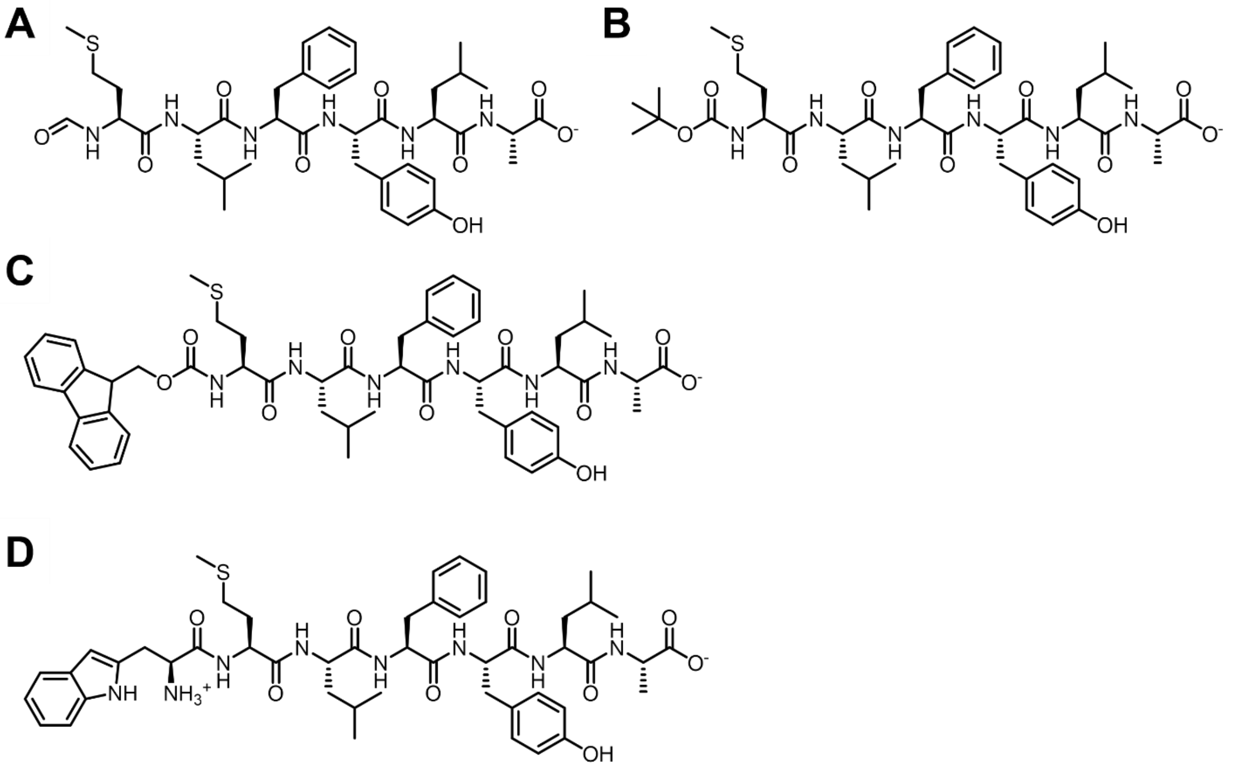

Figure S12: Structures of MLFYLA series (A) fMLFYLA (B) Boc-MLFYLA (C) Fmoc-MLFYLA (D) Trp-MLFYLA.

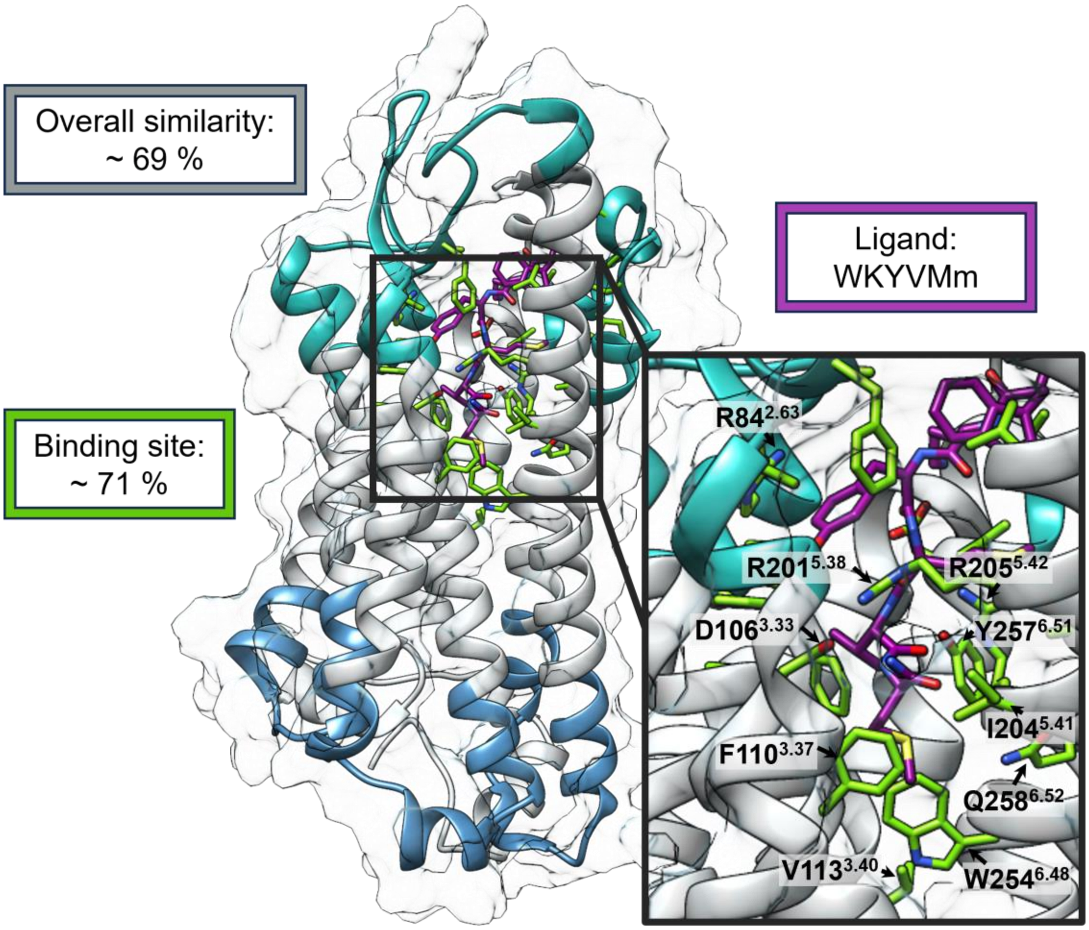

Figure S13: FPR1 homology model constructed with the FPR2 crystal structure (PDB ID: 6LW5[8]). The overall structural and binding-site similarity between FPR1 and FPR2 is shown. Binding-site residues are defined as residues within 4.5 Å of the ligand WKYMVm. The extracellular region is coloured cyan and the intracellular region blue.

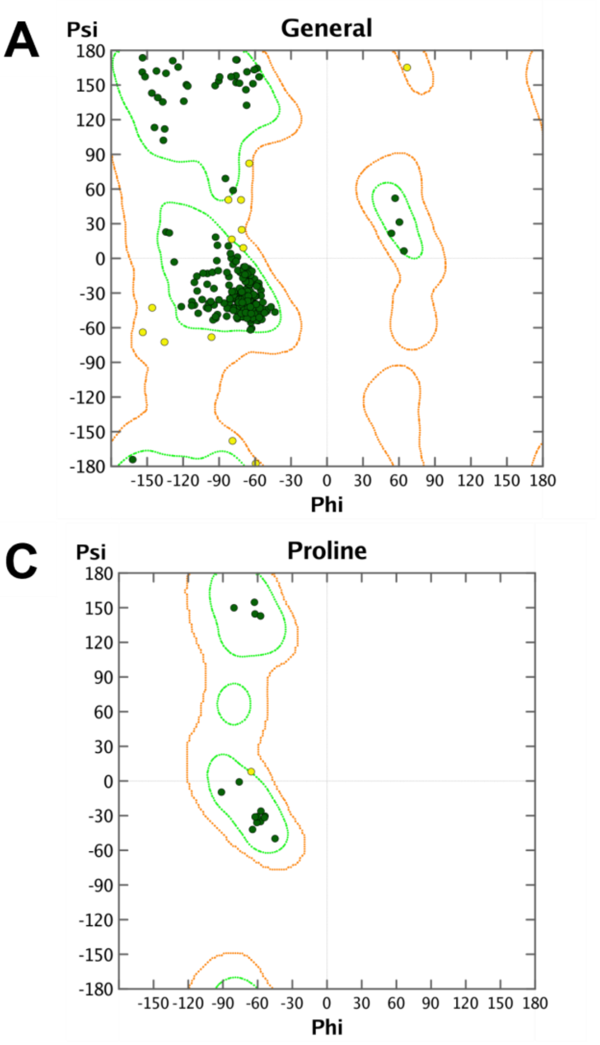

Figure S14: Ramchandran plots of the homology models. Green dots signify core values and yellow dots are allowed values.

**Synthesis of Fmoc-MLF and Fmoc-FLFLF**

**HPLC Method for Purity**

- Pump: Dionex UltiMate 3000
- Autosampler: Dionex UltiMate 3000
- Detector: Dionex UltiMate 3000
- Detection Wavelength: 210 nm
- Column: LiChrospher® 60 RP –select B (5 µm), 250-4 mm cartridge
- Solvent A: $H_2O$ with 0.05 % (*v*/*v*) trifluoroacetic acid
- Solvent B: $CH_3CN$ with 0.05 % (*v*/*v*) trifluoroacetic acid
- Gradient: (A %) 0 - 4 min: 90 %, 4 - 29 min: gradient 90 % to 0 %, 29 - 31 min: 0 %, 31 - 32 min: gradient 0 % to 90 %, 32 - 40 min: 90 %.
- Flow Rate: 1.00 mL/min
- Injection volume: 5 µL
- Room temperature

***N*-{[(9H-Fluoren-9-yl)methoxy]carbonyl}-(*S*)-methionyl-(*S*)-leucyl-(*S*)-phenylalanine (1)**

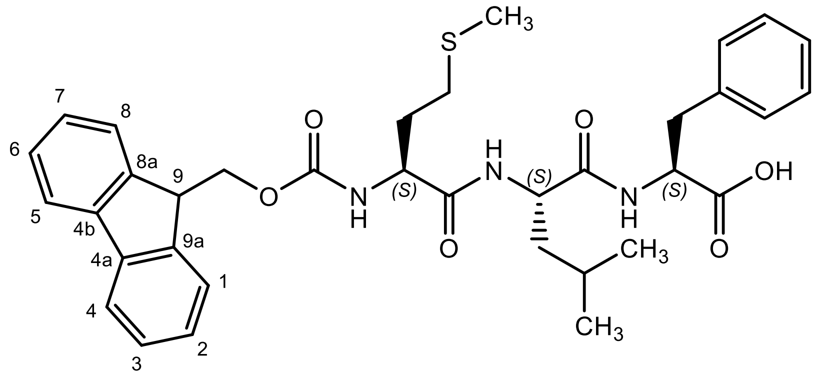

**1** was synthesized and purified at the Institute of Pharmaceutical and Medicinal Chemistry, University of Münster, Germany by Thomas Bödeker, member of the working group of Prof. Bernhard Wünsch. The zwitterionic tripeptide lithium (S)-methionyl-(S)-leucyl-(S)-phenylalaninate (35 mg, 84 µmol, 1.0 eq.) was added to a mixture of an aqueous $Na_2CO_3$ solution (10%, 1.5 mL) and 1,4-Dioxane (0.8 ml). At 0 °C, a solution of Fmoc-Cl (24 mg, 93 µmol, 1.1 eq.) in 1,4-Dioxane (1 mL) was added dropwise to the reaction mixture over 30 min under stirring. The reaction mixture was slowly warmed up to rt and stirred for approx. 1 h. The 1,4-Dioxane was evaporated *in vacuo* and the residue was diluted with $H_2O$ (10 mL). After acidification with HCl to pH ≈ 2, the solution was extracted with $CH_2Cl_2$ (4 x 10 mL). The organic layers were combined, dried ($Na_2SO_4$), filtered and the solvent was evaporated *in vacuo*. The crude product was purified by automatic flash chromatography; column 1: (Biotage® SNAP KP-Sil, 10 g, $CH_2Cl_2$:MeOH 100:0 → 93:7).

Colorless solid, **mp** 182 - 185 °C, **yield** 42 mg (79 %), $C_{35}H_{41}N_3O_6S$ ($M_r$ = 631.8), $\mathbf{R_f}$ = 0.41 (EtOAc:MeOH 4:1).

**$^1$H NMR** (600 MHz, DMSO-d$_6$): δ (ppm) = 0.80 (d, *J* = 6.6 Hz, 3H, CH(C*H*$_3$)$_2$), 0.84 (d, *J* = 6.6 Hz, 3H, CH(C*H*$_3$)$_2$), 1.35 – 1.44 (m, 2H, C*H*$_2$CH(CH$_3$)$_2$), 1.57 (nonett, *J* = 6.7 Hz, 1H, C*H*(CH$_3$)$_2$), 1.71 – 1.81 (m, 1H, SCH$_2$C*H*$_2$), 1.81 – 1.89 (m, 1H, SCH$_2$C*H*$_2$), 2.02 (s, 2.70H, SC*H*$_3$), 2.05* (s, 0.30H, SC*H*$_3$), 2.36 – 2.48 (m, 2H, SC*H*$_2$), 2.91 (dd, *J* = 13.9/8.1 Hz, 1H, PhC*H*$_2$), 3.05 (dd, *J* = 13.9/5.3 Hz, 1H, PhC*H*$_2$), 4.08 (td, *J* = 8.9/4.8 Hz, 1H, C*H*$_{(methionine)}$), 4.22 (t, *J* = 6.7 Hz, 1H, 9-*H*$_{(fluorene)}$), 4.24 – 4.32 (m, 3H, C*H*$_2$OCO, C*H*$_{(leucine)}$), 4.32 – 4.39 (m, 1H, C*H*$_{(phenylalanine)}$), 7.10 – 7.21 (m, 3H, 2-*H*$_{(Ph)}$, 4-*H*$_{(Ph)}$, 6-*H*$_{(Ph)}$), 7.21 – 7.26 (m, 2H, 3-*H*$_{(Ph)}$, 5-*H*$_{(Ph)}$), 7.32 (td, *J* = 7.5/1.1 Hz, 2H, 2-*H*$_{(fluorene)}$, 7-*H*$_{(fluorene)}$), 7.41 (t, *J* = 7.5 Hz, 2H, 3-*H*$_{(fluorene)}$, 6-*H*$_{(fluorene)}$), 7.56 (d, *J* = 8.3 Hz, 1H, N*H*CO$_2$), 7.71 (dd, *J* = 7.5/1.1 Hz, 2H, 1-*H*$_{(fluorene)}$, 8-*H*$_{(fluorene)}$), 7.89 (d, *J* = 7.5 Hz, 2H, 4-*H*$_{(fluorene)}$, 5-*H*$_{(fluorene)}$), 7.91 (d, *J* = 8.4 Hz, 1H, N*H*$_{(leucine)}$), 7.94 – 8.08 (m, 1H, N*H*$_{(phenylalanine)}$), 12.20 (bs, 1H, CO$_2$*H*).
The signal of the minor rotamer is marked with *. The ratio of the rotamers is 90:10.

**$^{13}$C NMR** (151 MHz, DMSO-d$_6$): δ (ppm) = 14.7 (1C, S*C*H$_3$), 21.7 (1C, CH(*C*H$_3$)$_2$), 23.0 (1C, CH(*C*H$_3$)$_2$), 24.0 (1C, *C*H(CH$_3$)$_2$), 29.7 (1C, S*C*H$_2$), 31.7 (1C, SCH$_2$*C*H$_2$), 36.4 (1C, Ph*C*H$_2$), 41.0 (1C, *C*H$_2$CH(CH$_3$)$_2$), 46.7 (1C, *C*-9$_{(fluorene)}$), 50.9 (1C, *C*H$_{(leucine)}$), 53.5 (1C, *C*H$_{(phenylalanine)}$), 53.8 (1C, *C*H$_{(methionine)}$), 65.6 (1C, *C*H$_2$CO$_2$), 120.1 (2C, *C*-4$_{(fluorene)}$, *C*-5$_{(fluorene)}$), 125.3 (2C, *C*-1$_{(fluorene)}$, *C*-8$_{(fluorene)}$), 126.3 (1C, *C*-4$_{(Ph)}$), 127.0 (2C, *C*-2$_{(fluorene)}$, *C*-7$_{(fluorene)}$), 127.6 (2C, *C*-3$_{(fluorene)}$, *C*-6$_{(fluorene)}$), 128.0 (2C, *C*-3$_{(Ph)}$, *C*-5$_{(Ph)}$), 129.1 (2C, *C*-2$_{(Ph)}$, *C*-6$_{(Ph)}$), 137.7 (1C, *C*-1$_{(Ph)}$), 140.7 (2C, *C*-4a$_{(fluorene)}$, *C*-4b$_{(fluorene)}$), 143.7 (1C, *C*-9a$_{(fluorene)}$), 143.9 (1C, *C*-8a$_{(fluorene)}$), 155.9 (1C, CH$_2$*C*O$_2$), 171.1 (1C, *C*=O$_{(methionine)}$), 171.6 (1C, *C*=O$_{(leucine)}$), 172.7 (1C, *C*O$_2$H).

**Purity (HPLC method):** purity = 99.2 %, $t_R$ = 21.4 min.

**IR (ATR):** ṽ [$cm^{-1}$] = 3290 (N-H), 2970 (C-H $_{(aliph)}$), 1690 (C=O $_{(carboxylic\ acid)}$), 1641 (C=O $_{(amide)}$), 1531 (C=C $_{(arom)}$).
**HRMS (APCI):** m/z = 632.2840 (calcd. 632.2789 for $C_{35}H_{42}N_3O_6S$ $[M+H]^+$).

**{[(9H-Fluoren-9-yl)methoxy]carbonyl}-(*S*)-phenylalanyl-(*S*)-leucyl-(*S*)-phenylalanyl-(*S*)-leucyl-(*S*)-phenylalanine (2)**

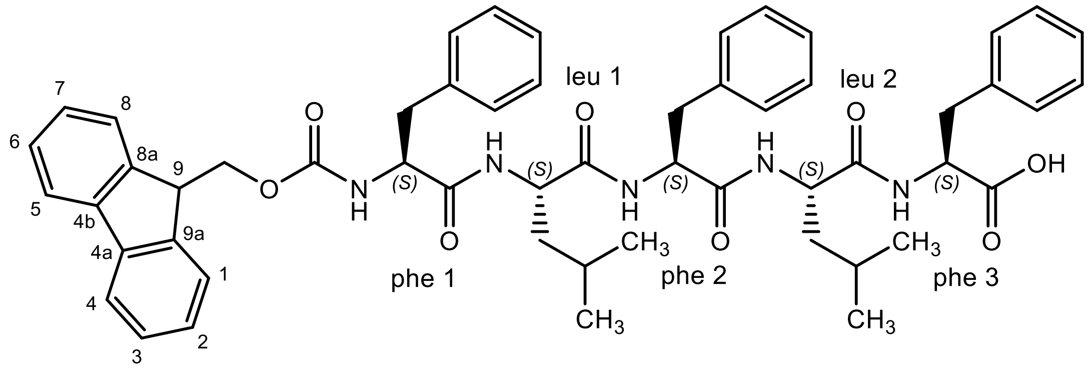

**2** was synthesized and purified in the Center for Soft Nanoscience in Münster, Germany by Luca Burg, member of the working group of Prof. Dr. Bart Jan Ravoo. A solid phase peptide synthesizer (SPPS) of the model *CS136XT* made by CSBio (Menlo Park, CA, USA) was used. The amino acids were bound to a 2-chlorotrityl chloride resin. *N*,*N'*-diisopropylcarbodiimide (DIC) / 1-hydroxybenzotriazole (HOBt) (0.4 M solution in DMF) was used as coupling reagents. DMF were used as solvent. The deprotection was performed with piperidine (20 % solution in DMF). The equivalents are relative to the active functionalities on the resin.
Fmoc-(*S*)-phenylalanine (1.5 eq.) was dissolved in dry $CH_2Cl_2$ (20 mL). The solution was added to the resin (1.6 mmol/g) under Ar. DIEA (2 eq.) was added and the mixture was agitated for 5 min by the argon stream. A second portion of DIEA (3 eq.) was added. After being agitated for 2 h by the Ar, MeOH (1 mL/g resin) was added and the resulting mixture was agitated for further 15 min to quench the remaining resin functionalities. The resin was filtered off and washed with $CH_2Cl_2$ (3 x 20 mL), DMF (3 x 20 mL), $CH_2Cl_2$ (3 x 20 ml) and MeOH (3 x 20 mL). The resin was dried *in vacuo* to determine the loading ratio by the weight increase. After the couplings were carried out by the SPPS, the peptide was cleaved off the resin and purified.
The resin was suspended in a solution of TFA:$H_2O$:Triisopropylsilane (95:2.5:2.5, 20 ml) and the mixture was stirred for 4 h. The resin was filtered off and was washed with TFA (3 x 5 mL). The peptide was precipitated by addition of a cold $Et_2O$/pentane solution (3:1). The precipitate was collected by centrifugation and the remaining water was removed by lyophilization to obtain **26**.
Colorless solid, **mp** 228 - 231 °C (decomposition), $C_{54}H_{61}N_5O_8$ ($M_r$ = 908.1), $\mathbf{R_f}$ = 0.36 (EtOAc:MeOH 4:1).
**$^1$H NMR** (600 MHz, DMSO-$d_6$): δ (ppm) = 0.76 – 0.87 (m, 12H, CH(C*H*$_3$)$_{2(leu\ 1,\ leu\ 2)}$), 1.38 (ddd, *J* = 7.6/7.3/3.0 Hz, 4H, C*H*$_2$CH(CH$_3$)$_{2(leu\ 1,\ leu\ 2)}$), 1.54 (dsept, *J* = 13.3/6.7 Hz, 2H, C*H*(CH$_3$)$_{2(leu\ 1,\ leu\ 2)}$), 2.72 (dd, *J* = 13.8/11.0 Hz, 1H, C*H*$_2$Ph$_{(phe\ 1)}$), 2.77 (dd, *J* = 14.1/9.0 Hz, 1H, C*H*$_2$Ph$_{(phe\ 2)}$), 2.88 – 2.95 (m, 2H, C*H*$_2$Ph$_{(phe\ 1,\ phe\ 3)}$), 2.98 (dd, *J* = 14.1/4.7 Hz, 1H, C*H*$_2$Ph$_{(phe\ 2)}$), 3.05 (dd, *J* = 14.0/5.3 Hz, 1H, C*H*$_2$Ph$_{(phe\ 3)}$), 4.09 – 4.18 (m, 3H, C*H*$_2$OCO, 9-*H*$_{(fluorene)}$), 4.22 – 4.32 (m, 2H, C*H*$_{(phe\ 1,\ leu\ 1)}$), 4.34 (q, *J* = 8.0 Hz, 1H, C*H*$_{(leu\ 2)}$), 4.43 (ddd, *J* = 8.6/7.7/5.4 Hz, 1H, C*H*$_{(phe\ 3)}$), 4.54 (td, *J* = 8.5/4.7 Hz, 1H, C*H*$_{(phe\ 2)}$), 7.04 – 7.36 (m, 17H, 2 – 6-*H*$_{(Ph,\ phe\ 1,\ phe\ 2,\ phe\ 3)}$, 2-*H*$_{(fluorene)}$, 7-*H*$_{(fluorene)}$), 7.40 (tdd, *J* = 7.5/3.5/1.1 Hz, 2H, 3-*H*$_{(fluorene)}$, 6-*H*$_{(fluorene)}$), 7.57 (d, *J* = 8.8 Hz, 1H, N*H*$_{(phe\ 1)}$), 7.61 (dd, *J* = 9.4/7.8 Hz, 2H, 1-*H*$_{(fluorene)}$, 8-*H*$_{(fluorene)}$), 7.87 (d, *J* = 7.6 Hz, 2H, 4-*H*$_{(fluorene)}$,

5-*H*$_{(fluorene)}$), 7.95 (t, *J* = 8.4 Hz, 2H, N*H*$_{(phe\ 2,\ leu\ 2)}$), 8.04 (d, *J* = 8.3 Hz, 1H, N*H*$_{(leu\ 1)}$), 8.11 (d, *J* = 7.7 Hz, 1H, N*H*$_{(phe\ 3)}$), 12.68 (bs, 1H, $CO_2$*H*).

**$^{13}$C NMR** (151 MHz, DMSO-$d_6$): δ (ppm) = 21.6 (1C, CH(*C*$H_3$)$_{2(leu\ 2)}$), 21.7 (1C, CH(*C*$H_3$)$_{2(leu\ 2)}$), 23.0 (1C, CH(*C*$H_3$)$_{2(leu\ 1)}$), 23.1(1C, CH(*C*$H_3$)$_{2(leu\ 1)}$), 23.9 (1C, *C*H($CH_3$)$_{2(leu\ 2)}$), 24.0 (1C, *C*H($CH_3$)$_{2(leu\ 1)}$), 36.6 (1C, *C*$H_2$Ph$_{(phe\ 3)}$), 37.2 (1C, *C*$H_2$Ph$_{(phe\ 2)}$), 37.3 (1C, *C*$H_2$Ph$_{(phe\ 1)}$), 41.0 (1C, *C*$H_2$CH($CH_3$)$_{2(leu\ 1)}$), 41.2 (1C, *C*$H_2$CH($CH_3$)$_{2(leu\ 2)}$), 46.5 (1C, *C*-9$_{(fluorene)}$), 50.7 (1C, *C*H$_{(leu\ 2)}$), 51.1 (1C, *C*H$_{(leu\ 1)}$), 53.2 (2C, *C*H$_{(phe\ 2,\ phe\ 3)}$), 55.9 (1C, *C*H$_{(phe\ 1)}$), 65.6 (1C, *C*$H_2CO_2$), 120.1 (2C, *C*-4$_{(fluorene)}$, *C*-5$_{(fluorene)}$), 125.2 (2C, *C*-1$_{(fluorene)}$, *C*-8$_{(fluorene)}$), 126.1 (2C, *C*-4$_{(Ph,\ phe\ 2,\ phe\ 3)}$), 126.4 (1C, *C*-4$_{(Ph,\ phe\ 1)}$), 127.0 (2C, *C*-2$_{(fluorene)}$, *C*-7$_{(fluorene)}$), 127.6 (2C, *C*-3$_{(fluorene)}$, *C*-6$_{(fluorene)}$), 127.9 (4C, *C*-3$_{(Ph,\ phe\ 2,\ phe\ 3)}$, *C*-5$_{(Ph,\ phe\ 2,\ phe\ 3)}$), 128.1 (2C, *C*-3$_{(Ph,\ phe\ 1)}$, *C*-5$_{(Ph,\ phe\ 1)}$), 129.0 (2C, *C*-2$_{(Ph,\ phe\ 3)}$, *C*-6$_{(Ph,\ phe\ 3)}$), 129.1 (2C, *C*-2$_{(Ph,\ phe\ 2)}$, *C*-6$_{(Ph,\ phe\ 2)}$), 129.2 (2C, *C*-2$_{(Ph,\ phe\ 1)}$, *C*-6$_{(Ph,\ phe\ 1)}$), 137.4 (1C, *C*-1$_{(Ph,\ phe\ 3)}$), 137.6 (1C, *C*-1$_{(Ph,\ phe\ 2)}$), 138.2 (1C, *C*-1$_{(Ph,\ phe\ 1)}$), 140.6 (2C, *C*-4a$_{(fluorene)}$, *C*-4b$_{(fluorene)}$), 143.7 (2C, *C*-8a$_{(fluorene)}$, *C*-9a$_{(fluorene)}$), 155.7 (1C, $CH_2$*C*$O_2$), 170.4 (1C, *C*=O$_{(phe\ 2)}$), 171.3 (1C, *C*=O$_{(phe\ 1)}$), 171.7 (2C, *C*=O$_{(leu\ 1)}$, *C*=O$_{(leu\ 2)}$), 172.7 (1C, *C*$O_2$H).

**Purity (HPLC method):** purity = 95.5 %, $t_R$ = 23.9 min.

**IR (ATR):** ṽ [$cm^{-1}$] = 3275 (N-H), 3080 (C-H $_{(arom)}$), 1697 (C=O $_{(carbamate)}$), 1635 (C=O $_{(amide)}$), 1539 (C=C $_{(arom)}$).

**MS:** m/z = 930.44 (calcd. 930.4412 for $C_{54}H_{61}N_5O_8$ $[M+H+Na]^+$).
